# Systematic Errors in DC Offset Measurement and Mitigation at Neurostimulation Electrodes: Causes, Implications and Solutions

**DOI:** 10.64898/2025.12.08.691908

**Authors:** Aaron K Gholston, Ricardo A Sanchez, Ty B Bigger, Spencer R Averbeck, Loren Rieth, Kip A Ludwig, James K Trevathan

## Abstract

**Objective:** Neuromodulation therapies are becoming increasingly common as a therapeutic strategy to treat neurological disorders and deficits. Despite their widespread use, the frameworks used to evaluate device longevity and electrochemical safety are limited, producing variable results for platinum (Pt) electrodes across studies. In addition, next-generation neuromodulation electrodes like thin-film microelectrode arrays (TFMEAs) are more prone to failure by virtue of their smaller size, higher edge to bulk ratios, and greater sensitivity to electrochemical dissolution processes. Two potential contributors to the variability in reported electrochemical safety limits and device performance are particularly sensitive to scaling electrodes to smaller geometries, namely: 1) insufficient measurement device input impedance and 2) unmitigated stimulator leakage currents.

**Approach:** To explore these effects, we first characterize the impact of electrode size (1.27 × 10^-4^ to 7.85 × 10^-3^ cm^2^) on apparent open circuit potential (V_aOCP_) and interpulse potential (V_IPP_) measurements at Pt electrodes across measurement instrumentation configurations. Next, we evaluated the effectiveness of commonly used DC mitigation strategies, including stimulation with capacitive coupling (CC), and with capacitive coupling combined with cathode/anode shorting after each pulse (CC+ES), as a function of electrode size.

**Main Results:** Without proper DC mitigation, insufficient measurement input impedance leads to an underestimation of DC offsets (ΔV_IPP_) and, consequently, a misrepresentation of the charge injection limit (Q_inj_) for Pt electrodes. In contrast, when DC offsets are properly mitigated and the input impedance of the measurement device is sufficient, the experimentally derived charge injection limits for Pt electrodes are larger than what has been reported historically.

**Significance:** By identifying the instrumentation-related confounds biasing measurements of electrochemical safety limits, this work provides a more consistent basis for the assessment of Q_inj_ for safety, as well as a framework for identifying and mitigating DC offset as a potential source of variability in electrophysiological findings across studies using TFMEAs.

## 1. Introduction

Implantable devices to stimulate the nervous system, also known as neuromodulation, bioelectronic medicine or electroceutical devices, are a staple of basic neuroscience studies and an increasingly common clinical therapy. The neuromodulation industry, including diverse therapies like cochlear implants for hearing restoration and deep brain stimulation (DBS) for Parkinson’s disease, is projected to exceed $10 billion in yearly revenue by 2030 [1]. With the implanted patient population increasing rapidly, designing neuromodulation devices that maximize therapeutic efficacy while ensuring device longevity remains a critical challenge.

For decades, platinum (Pt) and platinum alloys have served as the dominant electrode materials for neural stimulation devices due to their chemical inertness and corrosion tolerance. Nonetheless, these materials are not immune to degradation. Within the cochlear implant literature, a number of post-mortem analyses have revealed Pt deposits in adjacent tissue, indicating that Pt macroelectrodes undergo measurable corrosion under clinically relevant charge densities [2,3]. Historically, these findings have been dismissed as clinically insignificant because they have not been linked to adverse health outcomes or persistent device failures [4]. Nevertheless, as devices become smaller, the amount of dissolution that is tolerable before catastrophic device failure decreases drastically.

Efforts to reduce surgical invasiveness and improve the spatial precision of neural activation have led to the development and use of thin-film microelectrode arrays (TFMEAs) in both preclinical and clinical neuromodulation devices. These advancements have highlighted a unique limitation of TFMEAs: their susceptibility to corrosion-related failures compared to traditional macroelectrodes [5,6]. This vulnerability arises from multiple effects inherent to miniaturization. For a given applied current, smaller electrodes produce much higher charge densities, which increase the likelihood of undesirable electrochemical reactions associated with water electrolysis and corrosion. In addition, TFMEAs have limited material volume and higher edge-to-bulk ratios, making them more vulnerable to premature failure than macroelectrodes [5–7]. Despite numerous reports of premature failure of TFMEAs, the precise mechanisms underlying these failures remain poorly understood. Additionally, the safety frameworks used to define safe stimulation parameters were developed using Pt macroelectrodes and do not account well for the increased electrochemical vulnerability of TFMEAs [8–11].

The current frameworks used to evaluate the electrochemical and biological safety of neuromodulation devices were either derived from in-vitro voltage transients (VTs) [12] or phenomenological observations from in-vivo animal studies [10,13]. These frameworks have resulted in widely variable safety limits across studies ranging from 20-300 µC/cm^2^ for Pt macroelectrodes [12,14,15]. To adhere to these limits in practice, most clinical neuromodulation devices utilize charge-balanced, current-controlled waveforms to elicit neural activity while minimizing the risk of electrode corrosion and the accumulation of toxic byproducts [16]. However, the application of charge-balanced current waveforms can induce measurable shifts in the interpulse potential (V_IPP_) of the stimulation electrode, a phenomenon referred to as potential ratcheting, residual voltage, or the accumulation of a direct current (DC) offset [17–20]. The presence of significant DC offsets is known to accelerate electrode corrosion, alter local pH, and cause tissue damage [18,21–23]. Moreover, significant DC offsets may indicate the presence of unmitigated stimulator leakage currents, which themselves may activate or inhibit nervous tissue, resulting in altered therapeutic efficacy [21,24–28]. Importantly, unaccounted-for residual DC may represent an underrecognized source of experimental variability in preclinical neuromodulation studies. Therefore, ensuring device safety, efficacy and longevity requires both precise quantification and effective mitigation of DC offset accumulation.

Although the consequences of persistent DC offsets have been extensively studied, the influence of instrumentation on experimentally derived charge injection limits (Q_inj_) in the presence of these offsets has not been properly investigated. A methodological survey conducted herein revealed a number of studies reporting Q_inj_ from VT analysis which may be using measurement devices with insufficient input impedance (i.e. oscilloscopes) or failing to adequately minimize stimulator leakage currents. **We hypothesize that these instrumentation-related confounds have systematically misrepresented experimentally derived Q_inj_ thereby contributing to the wide variability in reported electrochemical safety limits across the neuromodulation literature**.

To evaluate the extent of these contributions, we first characterize the impact of electrode size (ranging from 1.27 × 10^-4^ to 7.85 × 10^-3^ cm^2^) on apparent open circuit potential (V_aOCP_) measurements at platinum electrodes across measurement instrumentation configurations. We demonstrate that the mismeasurement of the DC offsets due to probe-loading can dramatically influence experimentally determined limits for safe stimulation as assessed by voltage transient (VT) analysis, especially with progressively smaller electrodes. We then evaluate commonly used DC mitigation strategies, including stimulation with capacitive coupling, and with capacitive coupling combined with cathode/anode shorting after each pulse, as a function of electrode size. Collectively, these results show how DC offsets that develop on small electrodes are easily mismeasured when an oscilloscope is used. When appropriately measured, common DC mitigation strategies, like the use of a DC blocking capacitor, are often insufficient in isolation to completely mitigate stimulator leakage currents at small electrodes. Together, these data help explain the wide variation in reported safety limits, as well as the inconsistency in evoked electrophysiological responses observed across the literature [6,27,29–31]. Based on these results, we outline critical data that should be reported in every study, whether it be intended to assess safety for human use, or to improve rigor and reproducibility across fundamental neuroscience studies.

## 2. Methods

### 2.1 Electrode Fabrication and Polishing

90-10 platinum-iridium (Pt-Ir) disk electrodes of three different diameters; 1 mm, 0.250 mm and 0.127 mm (Surepure Chemicals PN:1884, Alfa Aesar PN:39383, AM Systems PN: 778000, respectively) with surface areas of 7.85 × 10^-3^, 4.91 × 10^-4^ and 1.27 × 10^-4^ cm^2^, were prepared. For each electrode, a 76.2 mm copper wire was soldered to 5 mm length of uncoated platinum wire. Soldered connections were cleaned via ultrasonic cleaner (Fisherbrand, PN:FB11201) at a frequency of 37 Hz and a power of 100% in DI water with a few drops of surfactant for 10 minutes. Post cleaning, the platinum electrode protrusion was encased in EPO TEK 301 (PN:FB194-ND) epoxy and allowed to cure. Once cured, electrodes were polished to 10 µm grit and cleaned once more ultrasonically using the same parameters. All electrodes were also polished and cleaned periodically between voltage transient experiments to standardize the surface condition of the electrodes [32].

### 2.2 Electrochemical Characterization

Electrochemical impedance spectroscopy (EIS) was performed on all samples. Additionally, a set of apparent open circuit potential (V_aOCP_) and voltage transient (VT) measurements were collected for each of the platinum electrodes (n = 12) at 4 different effective input impedances: 1 MΩ, 10 MΩ, 10 GΩ, and 10 TΩ. All electrochemical measurements were performed in 1X (0.01 M) phosphate buffered saline (P38135, Sigma-Aldrich) utilizing a 3-electrode setup against a large platinum counter electrode (Metrohm, 1 cm^2^) and a single junction Ag|AgCl reference electrode (BASi, PN: MF-2056) filled with 3M KCl. All electrochemical measurements were performed at room temperature. **(Figure 1(a))**

**Figure 1:**
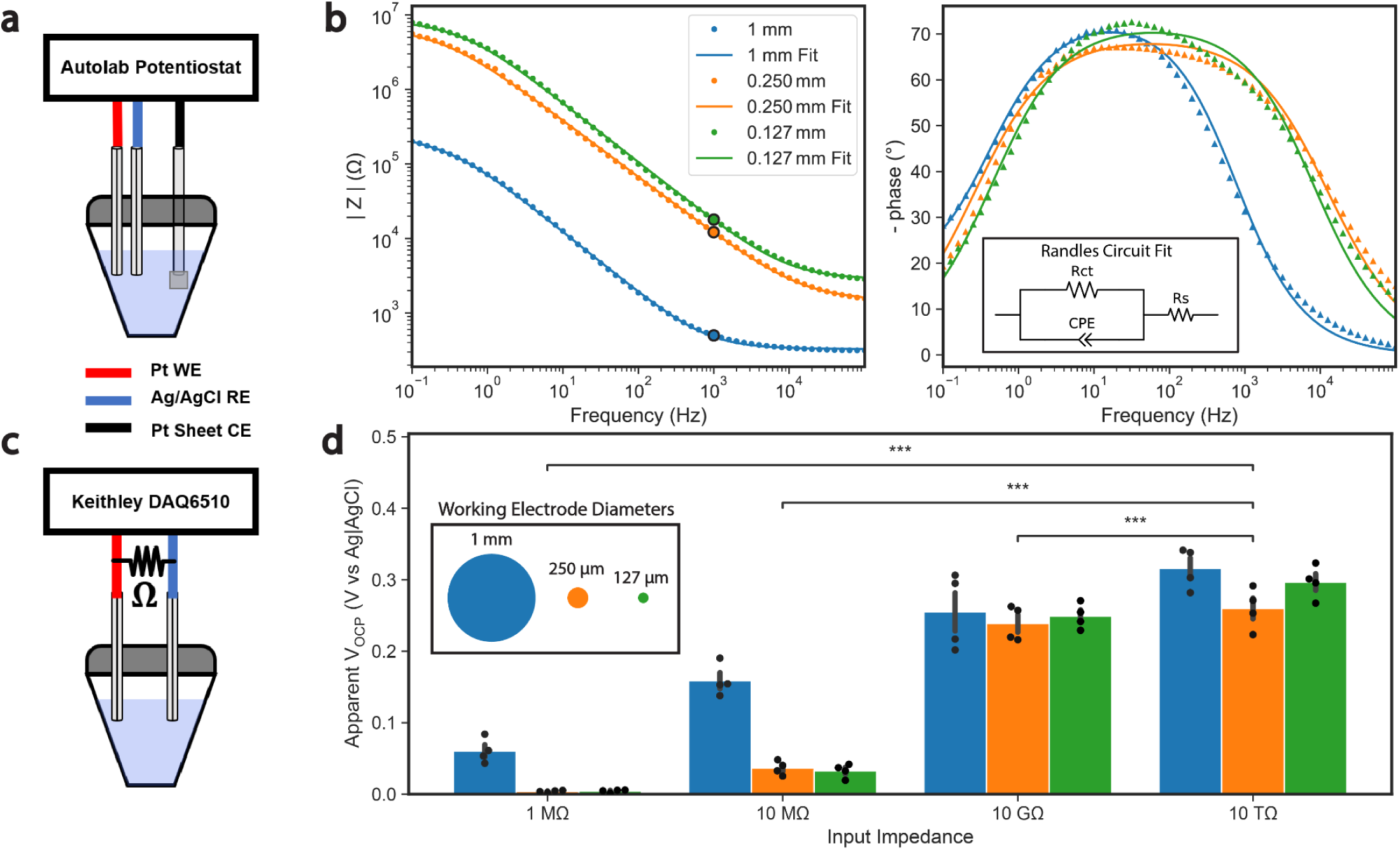
Characterization of DC electrode impedance (R_dc_) and DC electrode voltage (V_aOCP_) across input impedances. (a) Diagram of electrode configuration used to conduct EIS measurements. (b) Mean modeled vs experimental bode impedance magnitude and phase for electrodes of 1 mm (blue), 250 µm (orange) and 127 µm (green) diameter electrodes. Points outlined in black represent the impedance measured at 1 kHz. The Randles circuit diagram used to determine fit parameters from EIS spectra is present within the phase plot. (c) Diagram of electrode configuration used to conduct apparent open circuit potential (V_aOCP_) measurements. (d) Apparent OCPs measured at different input impedances across electrode sizes. Legend in panel d is a scaled representation of the disk electrode diameters. All data in panel d are represented as mean ± SEM. *** for p < 0.001.

#### 2.2.1 Electrochemical Impedance Spectroscopy (EIS) Measurements

EIS measurements were performed using an Autolab potentiostat (PG-STAT12, Metrohm) with a built-in frequency response analyzer (FRA2), EIS measurements were performed by applying 10 mV sine waves, at 0V and the open circuit potential (OCP) relative to a Ag|AgCl reference electrode. Impedance spectra were collected from 0.1 Hz to 100 kHz, spaced at 10 points per decade and subsequently fitted with an equivalent Randles circuit to determine charge transfer resistance (R_ct_), solution resistance (R_s_) and constant phase element (CPE) values. (**Figure 1(b), Supplementary Figure 1, Supplementary Table 1, Supplementary Table 2**)

#### 2.2.2 Apparent Open Circuit Potential Measurements at Equilibrium

For the V_aOCP_ measurements, electrodes of 1 mm (n=4), 0.25 mm (n=4) and 0.127 mm (n=4) diameter were placed in 1X PBS and connected to a Keithley DAQ6510 multimeter (**Figure 1(c)**). The three electrode sizes tested were selected based off pilot studies and were intended to span the range where we begin to observe appreciable probe loading effects using common oscilloscopes (i.e. typically with input impedances of 1-10 MΩ). The DAQ6510 multimeter was selected because it has a maximum input impedance greater than 10 GΩ as well as the ability to conduct measurements at 10 MΩ input impedance. 1 MΩ V_aOCP_ measurements were obtained by adding a 1MΩ oscilloscope probe in parallel with the DAQ6510. 10 TΩ V_aOCP_ measurements were obtained by using the guard port of a Keithley 6221 current source. The V_aOCP_ of each electrode was measured with respect to a Ag|AgCl electrode and allowed to stabilize until a 0.5 mV/s change in potential was observed. Three successive measurements were performed on each electrode and then averaged to produce the value presented in **Figure 1(d)**.

#### 2.2.3 Voltage Transient Measurements

VT measurements were performed at all input impedances by applying cathodic-leading, symmetric, charge-balanced biphasic waveforms (200 µs/phase, 50 µs interphase, 30 Hz, 750 pulses) using a Keithley 6221 current source. This current source was selected for its particularly high output impedance (>10 TΩ) which is essential to isolate the parasitics introduced strictly from the measurement instrumentation. Twenty logarithmically spaced charge densities between 2 and 600 µC/cm^2^ were applied to each electrode and the resulting voltage response was measured (**Supplementary Table 3**). These parameters were inspired by the seminal paper by Robblee, et al., presented in 1990 [12]. Electrodes were allowed to stabilize between trials until a 0.5 mV/s change in V_IPP_ was observed. The compliance voltage of the Keithley 6221 was set to 20V for all experiments. The interfacial electrode polarization (i.e. E_mc_ and E_ma_) were calculated on the last pulse of each charge density applied by subtracting the access voltage (V_a_) to isolate the polarization of the electrode-electrolyte interface with respect to Ag|AgCl [33,34]. The charge injection limit (Q_inj_) in this study is defined as the charge density in which either E_ma_ or E_mc_ exceeded 0.8 and -0.6 V, respectively. **(Figure 3(a))** Alternatively, in some studies, this value has also been referred to as the charge injection capacity [12,35]. It is recommended to perform a validation of the V_a_ subtraction to evaluate the performance of the Q_inj_ determination methodology using a secondary technique [36]. As validation, the calculated access voltages were converted into an equivalent value of R_s_ using the applied current. These values of R_s_ were then compared to the R_s_ values obtained from the EIS Randles fit (**Supplementary Table 4, Supplementary Figure 3**).

#### 2.2.4 Implementation of Common DC Mitigation Strategies

VT measurements were also collected at the input impedances of 1 MΩ, 10 MΩ, and 10 GΩ using our custom-built DC mitigation strategy which deploys two 1µF, low leakage current DC blocking capacitors (PN: 105PHC400K) in series with the stimulator and a high off-impedance switch (PN: DG417L) behind the capacitors and in parallel with the stimulator output. The parasitics of the DC mitigation circuitry were considered and differential trace impedances were characterized to ensure they did not impact experimental results (**see Supplementary Methods**). Voltage transient experiments were run using capacitive coupling (CC) only as well as capacitive coupling with electrode shorting (CC+ES) (**Supplementary Figure 2**). In the capacitive coupling with electrode shorting case, the working electrode and counter electrode were shorted together following the application of each current pulse to prevent significant DC accumulation at the electrode interface. Under the capacitive coupling only condition, pulse trains were delivered to the working electrode through the DC blocking capacitors, however the working and counter electrodes were shorted between trials rather than between pulses.

### 2.3 Statistical Analyses

Statistical analyses were performed using Python (v3.10.11) with the SciPy library (v1.10.1). Normality was assessed using the Shapiro-Wilk test for both V_aOCP_ and Q_inj_ datasets. Results indicated a mix of normally and non-normally distributed groups; therefore, non-parametric statistical tests were used throughout. All statistical tests were conducted with a significance level of α = 0.05. All electrochemical data are reported throughout the text as mean ± standard error of the mean (SEM).

Wilcoxon signed-rank tests were used to assess statistical differences in VaOCP and Qinj measurements across input impedances. The V_aOCP_ and Q_inj_ data without DC mitigation compared each group to the 10 TΩ condition. The Q_inj_ data with DC mitigation compared each group to the 10 GΩ condition. A Holm-Bonferroni correction was applied to all Wilcoxon tests.

### 2.4 Methodological Survey

A methodological survey was conducted to identify studies reporting charge injection limits/capacities derived from voltage transient measurements across the neuromodulation literature. A unified Boolean search strategy was deployed using the following search string input into Google Scholar: ("voltage transient" OR "voltage excursion") AND ("charge injection limit" OR "charge injection capacity") AND ("oscilloscope"). This search yielded 165 records. Articles were filtered for duplicates and were screened by three unique reviewers to determine that they met the inclusion criteria (**Supplementary Methods**). Ultimately, this search strategy revealed 77 unique results in which a predetermined set of parameters was extracted (**Supplementary Methods, Supplementary Table 5**). Comprehensive results from the methodological survey are presented in a table in the supplement.

## 3. Results

The results presented in this manuscript highlight how measurement device input impedance, electrode size and DC mitigation strategies influence the accurate measurement and accumulation of DC offsets (ΔV_IPP_). **Section 3.1** examines how electrode size and input impedance impacts the measurement of the V_aOCP_. Then, in **Section 3.2**, these findings are contextualized using the voltage divider equation. In **Section 3.3**, we present voltage transient measurements and corresponding Q_inj_ collected at the same input impedances tested in **Section 3.1**. After establishing the input impedance required for reducing/eliminating the mismeasurement of ΔV_IPP_, we use this set-up to evaluate the effectiveness of common DC mitigation strategies. The impact of input impedance and DC mitigation strategy on the calculated Q_inj_ is summarized in **Section 3.4**.

### 3.1 DC Electrode Impedance & Open Circuit Potential Characterization

In practice the input impedance of a measurement device should be much larger (typically by a factor of at least 100) than the impedance of the circuit being measured [37]. This minimizes oscilloscope probe-loading, preserving both the accuracy of the measurement and the underlying circuit behavior. At DC, the impedance of an electrode as modeled by a Randles circuit, is purely resistive and simplifies to R_dc_=R_ct_ + R_s_, where R_ct_ is the charge transfer resistance and R_s_ is the solution resistance. As a result, R_dc_ is usually the highest impedance of an electrode, as shown on the representative Bode plot in **Figure 1(b)** [26–28]. Therefore, the electrode DC impedance is the most relevant measure to assess appropriateness of the input impedance of the measurement device. Initially, to estimate R_dc_ of our electrodes, the value of the charge transfer and solution resistances were determined from the Randles circuit fit of the EIS spectra collected around 0V vs Ag|AgCl (**Figure 1(b)**, **Table 1**; R_ct_ = 2.41 × 10^5^ ± 1.93×10^3^, 6.98×10^6^ ± 9.35×10^5^, 8.70×10^6^ ± 7.30×10^5^ Ω, R_s_ = 3.26× 10^2^ ± 5.10, 1.46 × 10^3^ ± 3.25× 10^1^, 2.83 × 10^3^ ± 1.34× 10^1^ MΩ for 1 mm, 0.250 mm and 0.127 mm diameter electrodes, respectively). Complete Randles circuit fit parameters for each electrode are presented in **Supplementary Table 1**.

**Table 1:** Average Randles circuit parameters extracted from EIS around 0V vs Ag|AgCl and OCP across electrode sizes. Comparison of 0V and OCP EIS spectra across sizes is presented in Supplementary Figure 4.

| Electrode Diameter<br>(mm) | R<sub>ct</sub> (Ω) |  | R<sub>s</sub> (Ω) |  | Q (Ω<sup>-1</sup> sec<sup>a</sup>) |  | n |  |
| --- | --- | --- | --- | --- | --- | --- | --- | --- |
|  | 0V | OCP | 0V | OCP | 0V | OCP | 0V | OCP |
| 1 mm | 2.41E+05 | 2.77E+06 | 3.26E+02 | 3.26E+02 | 2.89E-06 | 2.52E-06 | 8.20E-01 | 8.35E-01 |
| 0.250 mm | 6.98E+06 | 6.01E+07 | 1.46E+03 | 1.48E+03 | 1.08E-07 | 1.15E-07 | 7.71E-01 | 7.81E-01 |
| 0.127 mm | 8.69E+06 | 9.79E+07 | 2.83E+03 | 2.93E+03 | 5.81E-08 | 7.29E-08 | 8.04E-01 | 8.33E-01 |

Reported values of open circuit potential (OCP) for platinum electrodes in 1X PBS, at pH 7.4, typically fall between 0.25V and 0.30V versus a saturated Ag|AgCl reference electrode [38–40]. **Figure 1(d)** displays the apparent steady-state OCP (V_aOCP_) obtained by measurement devices with input impedances of 1 MΩ, 10 MΩ, 10 GΩ or 10 TΩ, respectively. At the 1 MΩ input impedance, the V_aOCP_ measurement dramatically underestimates the OCP at all electrode sizes compared to what would be expected from literature, with the V_aOCP_ at the two smallest electrode sizes approaching zero (60.4 ± 8.65, 3.87 ± 0.715, and 4.37 ± 0.985 mV for 1 mm, 0.250 mm and 0.127 mm diameters, respectively). At 10 MΩ, the V_aOCP_ at all electrode sizes are notably less than would be expected from literature (159 ± 11.1, 36.6 ± 4.90, and 32.7 ± 4.82 mV for 1 mm, 0.250 mm and 0.127 mm diameters, respectively); however, as expected, the largest electrode size with the lowest R_dc_ is much closer to established literature values. Only by using the highest input impedance configurations (10 GΩ & 10 TΩ) do the measured V_aOCP_ values, across all electrode sizes, approach the expected range from literature. When compared to the V_aOCP_ values collected at 10 TΩ, statistically significant pairwise differences were observed at the 1 MΩ, 10 MΩ, 10 GΩ input impedances across sizes.

### 3.2 Impact of Input Impedance on Measured Electrode Potential

Equations 1 & 2 represent a simplistic theoretical calculation of expected V_aOCP_ by modeling the DC electrode impedance (R_dc_) and measurement device impedance (Z_in_) as a voltage divider, with equation 2 being the simplified form in terms of the ratio of R_dc_/Z_in_ [38].

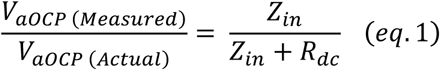

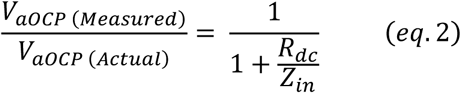

**Figure 2(a)** shows the relationship between increasing measurement device input impedance and R_dc_/Z_in_ for electrodes of decreasing surface area. The measured apparent OCP values were normalized to the values collected at the 10 TΩ input impedance for each sample and plotted against the R_dc_/Z_in_ (**Figure 2(b)**). The theoretical model defined by equations 1 and 2 is displayed as a black dashed line in **Figure 2(b)** and is overlaid with experimentally measured data to illustrate the relationship between the theoretical and observed V_aOCP_ values across varying R_dc_/Z_in_ ratios. As can be seen, the functional measurements approximate the theoretical calculation which generates a sigmoidal curve when plotted; however, this sigmoidal curve is shifted to the right for the functional measurements.

**Figure 2:**
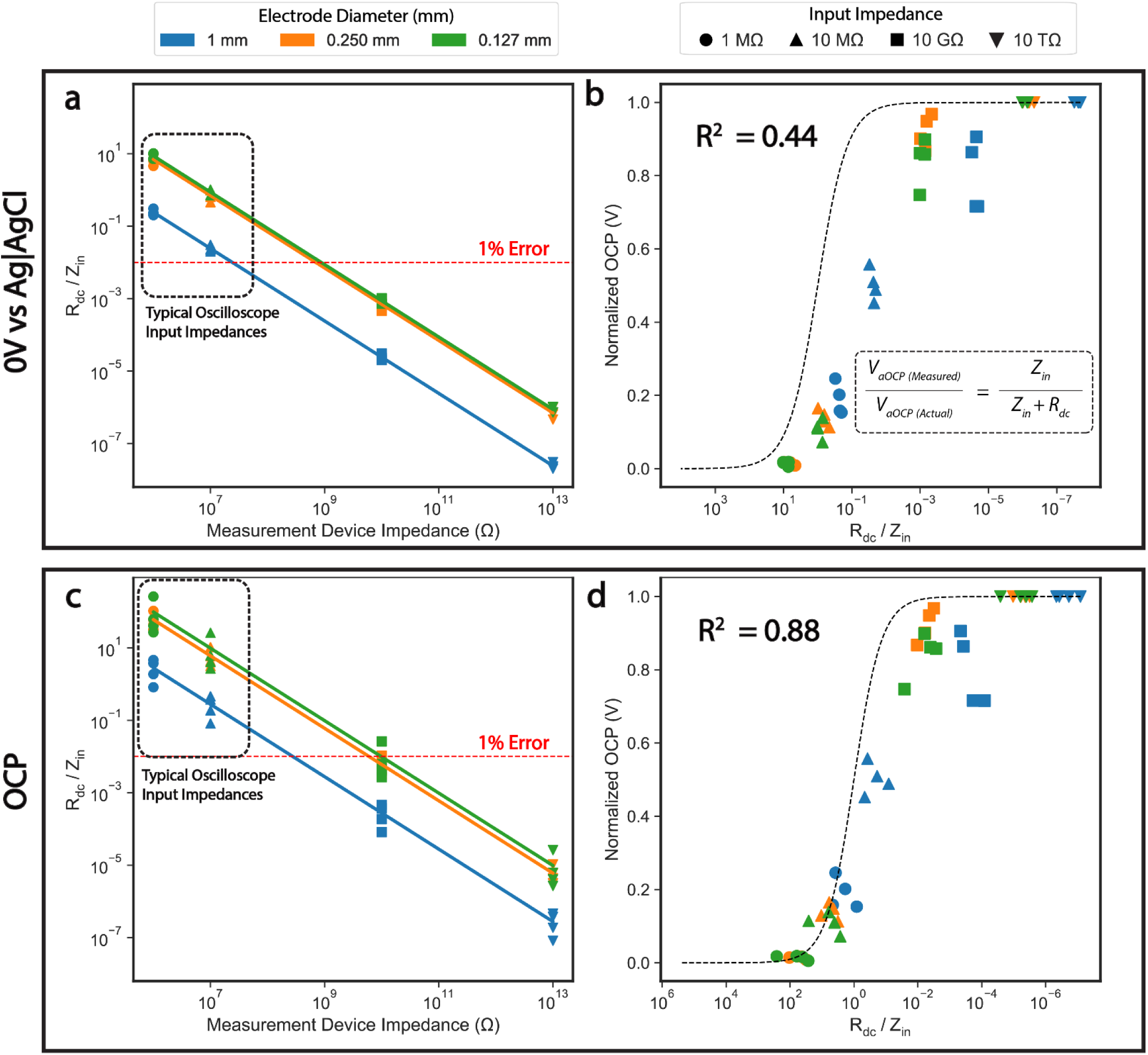
Significant measurement device backend impedance (Z_in_) relative to DC electrode impedance (R_dc_) is required to properly measure the DC electrode potential (V_aOCP_) of an electrode. (a) and (c) Measurement device impedance vs R_dc_ / Z_in_ for each electrode where R_dc_ is derived from the Randles circuit fit and defined as R_ct_ + R_s_ for Randles parameters taken from EIS performed around 0V vs Ag|AgCl and around OCP of each electrode, respectively. Dashed rectangle highlights typical input impedances for a standard oscilloscope probe. (b) and (d) R_dc_ / Z_in_ plotted against normalized apparent open circuit potential measurements. Black dashed line in both plots is the expected V_aOCP_ with the input impedance (Z_in_) and the DC electrode impedance (R_dc_) modeled as a voltage divider. Red dashed lines in panels a and c represent the ratio of R_dc_/Z_in_ in which the idealized voltage divider model reaches a 1% measurement error.

One possible source of this deviation of our functional measurements from theory is the method of EIS used to calculate the R_dc_ of each electrode. In our initial EIS measurements, we applied a sinusoid at each frequency centered on 0V vs Ag|AgCl. This is a common practice in neuromodulation for EIS measurements, because the OCP of an electrode is dependent on several factors (i.e. electrode surface state, concentration of redox species, etc.), creating a source of variability in the measurement of impedance at a specific frequency. Consequently, centering the sinusoids at 0V creates a common reference point mitigating this issue. However, applying a sinusoid centered at 0V forces the platinum electrodes away from their natural open circuit potential of ∼0.3V vs Ag|AgCl in 1X PBS. This requires inducing faradaic reactions, which in turn lowers R_ct_ [41,42].

To assess this possibility, we reconducted EIS with the applied sinusoids centered around the OCP of the electrodes; the OCP was measured for 60 seconds prior to the collection of each EIS spectrum. This should result in a much larger but more accurate estimate of R_ct_ **(Table 1)**. **Figure 2(c)** displays the new ratios of R_dc_/Z_in_ using this alternative approach. **Figure 2(d)** depicts the normalized OCP as a function of R_dc_/Z_in_ calculated using these new EIS values versus the same functional data presented in **Figure 2(b)**. The new calculated expectation of normalized OCP as a function of R_dc_/Z_in_ more closely approximates the measured functional data (R^2^ = 0.88 @ OCP versus R^2^ = 0.44 @ 0V vs Ag|AgCl), suggesting that performing EIS measures centered around the OCP of each electrode provided a more accurate representation of probe-loading effects.

### 3.3 Impact of R_dc_/Z_in_ on Apparent Charge Injection Limit

Voltage transients (VT) are commonly used to determine the safe limits of stimulation for an implanted electrode. By measuring electrode polarization during therapeutic stimulation pulses, VTs help identify conditions that may drive unwanted faradaic reactions, leading to electrode dissolution and/or the generation of toxic byproducts. For a platinum/platinum iridium electrode, a cathodic polarization of -0.6 volts or an anodic polarization of 0.8V relative to Ag|AgCl can cause the electrolysis of water, otherwise known as the ‘water window’ for platinum. This, in turn evolves H_2_ or O_2_, respectively, changing the pH of nearby tissue. Typically, the current/charge density just before these reactions occur is deemed the ‘charge injection limit’ (Q_inj_) [17,35]. In the context of voltage transient responses, it is important to understand the distinct components of the voltage waveform and what each represents. **Figure 3(a) and (b)** depict the experimental set-up for measuring VTs, identifies these distinct components and describes the process used in this study for determining the Q_inj_.

**Figure 3:**
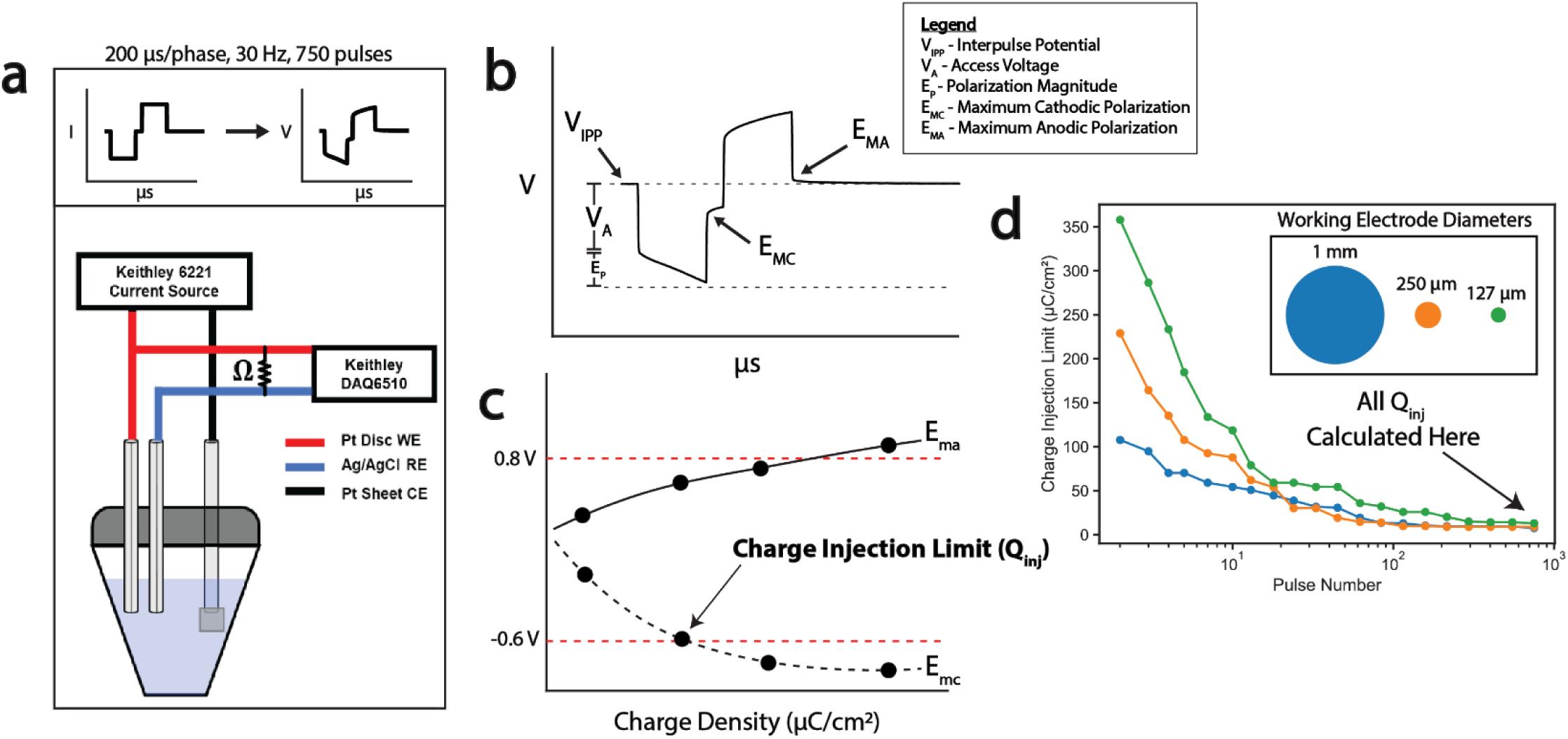
Experimental configuration and methodology used to determine charge injection limit (Q_inj_). (a) Instrumentation configuration and pulse parameters for voltage transient measurements. The top panel depicts representative input current and output voltage traces. (b) Representative voltage transient with different waveform components highlighted. (c) Diagram of E_mc_ and E_ma_ plotted as a function of charge density methodology for charge injection limit determination. (d) Theoretical charge injection limits calculated across pulses and sizes at 10 GΩ of input impedance.

In **Figure 3(b)**, V_IPP_ is defined as the interpulse potential, during which the stimulator current has been set to zero. In this study, V_IPP_ is used as a surrogate measure of DC electrode potential for our voltage transient experiments. Accurate measurement of the electrode’s interpulse potential (V_IPP_) during voltage transient experiments is crucial for determining when the water window is breached and, consequently, for establishing the Q_inj_ of a given electrode material. The access voltage (V_a_) arises from the instantaneous voltage drop across the electrolyte and is influenced by the electrolyte conductivity and cross-sectional area of the electrode. For biphasic stimulation waveforms, V_a_ appears as a step voltage change at the onset and termination of each phase due to the instantaneous change in current. V_a_ represents the voltage response due to the bulk impedance of the electrochemical system and is not associated with the electrochemical activity at the electrode interface; however, it is critical in the determination of the interfacial polarization voltage at the electrode.

In contrast, E_mc_ and E_ma_ represent the polarization experienced at the electrode interface in the cathodic and anodic direction, respectively. These values, relative to V_IPP_, provide a direct measure of the capacitive and faradaic processes occurring at the interface and are used to evaluate whether the electrode remains within safe electrochemical limits. The Q_inj_ in this study is defined as the charge density in which the E_mc_ or E_ma_ exceed the cathodic or anodic limit for the water window **(see** **Figure 3(c)).** As VTs require the application of many pulses before reaching stability, particularly those conducted without DC mitigation, all calculations of Q_inj_ were performed after 750 biphasic pulses were applied which reached a point of stability for all electrode sizes and charge densities tested (**see** **Figure 3(d)**) [43].

#### 3.3.1 Observed ΔV_IPP_ Increased with Sufficient Input Impedance

Given that one of the goals of this study is to evaluate how instrumentation input impedance influences the measurement of the DC offset (ΔV_IPP_), the data in this section are presented using a stimulation set-up anticipated to generate significant ΔV_IPP_, i.e. stimulation with charge-balanced biphasic pulses without the use of a DC blocking capacitor and shorting of the cathode and anode between pulses. **Figure 4(a)** depicts a representative voltage transient for each electrode size obtained as a function of measurement device input impedance. The charge density of ∼100 µC/cm^2^ was selected as it is the midpoint of the range of 50-150 µC/cm^2^, which has been identified as the Q_inj_ for Pt/Ir electrodes in prior studies. The largest ratio of electrode impedance to instrumentation impedance is at DC (i.e. R_dc_/Z_in_) thus V_IPP_ is anticipated to be the voltage transient component that is most impacted by changes in measurement device input impedance. Conversely, V_a_ is a very high frequency component of the VT. At this frequency the electrode impedance is dominated by R_s_ which is orders of magnitude lower than R_dc_. Consequently, the ratio of instrumentation impedance (Z_in_) to the high frequency electrode impedance (R_s_) for the measurement of V_a_ is orders of magnitude larger, therefore the measurement of V_a_ is anticipated to be minimally impacted by changes in measurement device input impedance.

**Figure 4:**
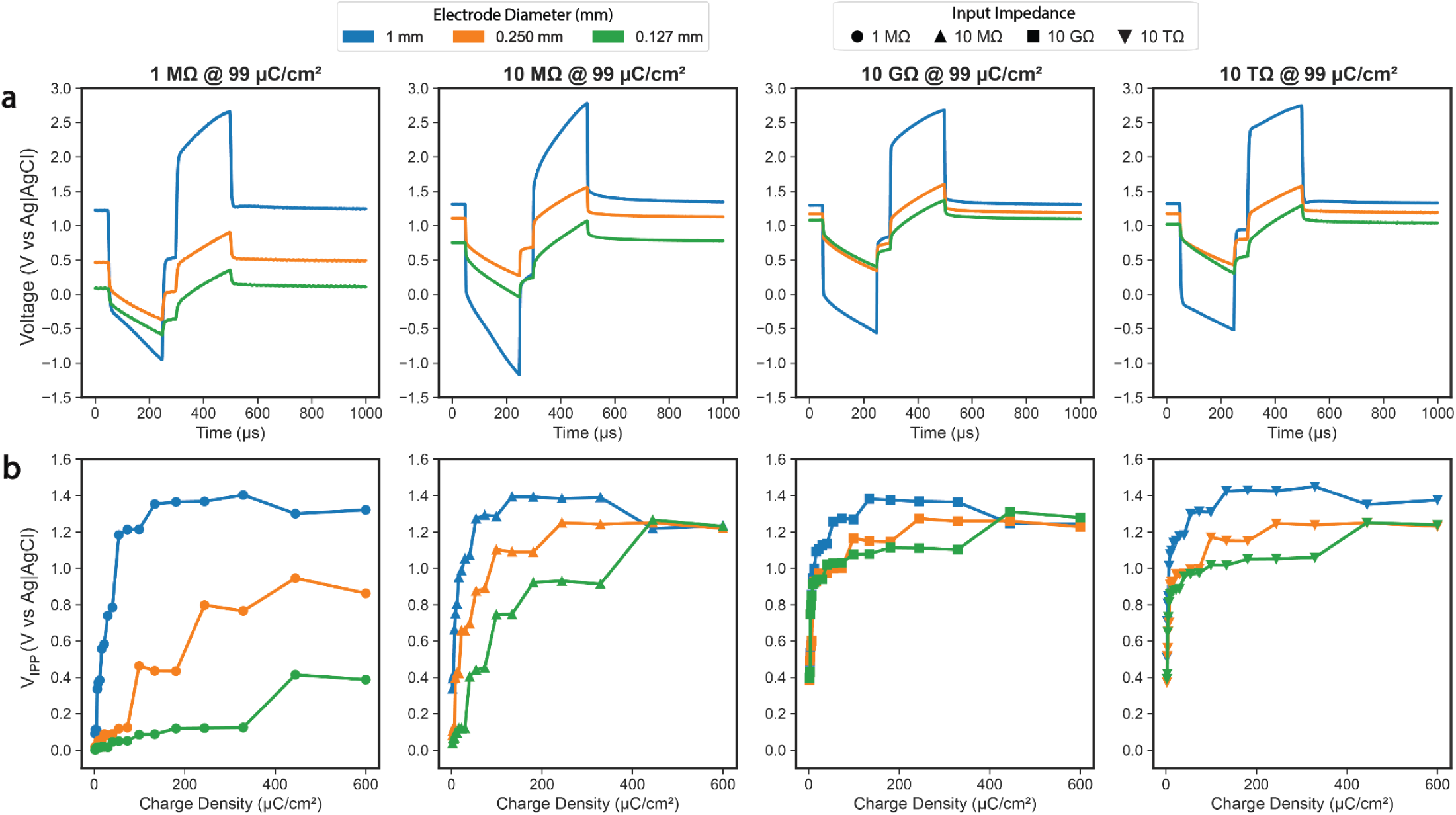
Observed changes in representative voltage transients and observed interpulse potential (V_IPP_) across input impedances. (a) Average representative voltage transients across sizes collected at ∼100 µC/cm^2^ across input impedances and electrode diameters. (b) Average V_IPP_ across input impedances and electrode diameters. Depicts the VT waveform shape and observed convergence of V_IPP_ at larger input impedances across increasingly smaller electrodes.

Consistent with theoretical expectations, as a measure of DC electrode potential, the measurement of V_IPP_ during VTs at each electrode size follows the same pattern as our prior measurements of V_aOCP_ (shown in **Figure 4(b))**. The measured V_IPP_ of the largest electrode size of 1 mm diameter which has the lowest DC impedance is the most consistent across the input impedances tested (an average of 1.21 ± 0.885 × 10^-2^, 1.28 ± 1.87 × 10^-2^, 1.27 ± 1.89 × 10^-2^ and 1.31 ± 1.73 × 10^-2^ V at 1 MΩ, 10 MΩ, 10 GΩ, and 10 TΩ, respectively). Conversely, the smallest electrode size of 0.127 mm diameter with a much higher DC impedance is most impacted by the input impedance of the measurement system (averaging 8.58 × 10^-2^ ± 1.45 × 10^-3^, 7.47 × 10^-1^ ± 1.65 × 10^-2^, 1.08 ± 0.510 × 10^-2^, and 1.02 ± 4.14 × 10^-3^ V at 1 MΩ, 10 MΩ, 10 GΩ, and 10 TΩ, respectively). Importantly, as the input impedance of the measurement device increases, the measured V_IPP_ at each electrode size converges to between 1.1 and 1.3V even at much greater charge densities than 100 µC/cm^2^ (see **Figure 4(b)**).

As expected, the measurements of V_a_ are relatively consistent across measurement system input impedances. To maintain constant charge density across electrode sizes, the applied current was scaled proportionally to the electrode surface area. Because the applied current scales with the square of the radius, the current delivered to the 1 mm diameter electrode was ∼61 times greater than that applied to the 0.127 mm diameter electrode. According to Ohm’s law (V_a_= I × R_s_) and based on the geometric relationship between R_s_ and electrode radius for circular electrodes described in equation 3, where R_s_ is the solution resistance, *ρ* is the electrolyte resistivity and r is the electrode radius. This difference in current and R_s_ results in an expected ∼8x larger V_a_ for the 1 mm diameter electrode compared to the 0.127 mm diameter electrode [33–35]. Across measurement input impedances of 1 MΩ, 10 MΩ, 10 GΩ and 10 TΩ, the average V_a_ measured during the VT for the 0.127 mm diameter electrode was 1.65 × 10^-1^ ± 3.03 × 10^-3^, 2.09 × 10^-1^ ± 1.11 × 10^-2^, 2.04 × 10^-1^ ± 6.31 × 10^-3^, and 1.88 × 10^-1^ ± 3.28 × 10^-3^ V, respectively. By comparison, the measured V_a_ during the VT for the 1 mm diameter electrode are ∼6-8x larger; 1.27 ± 6.01 × 10^-2^, 1.32 ± 2.96 × 10^-2^, 1.32 ± 3.20 × 10^-2^ and 1.34 ± 1.08 × 10^-1^ V for the 1 MΩ, 10 MΩ, 10 GΩ, and 10 TΩ measurement input impedances, respectively. The V_a_ for the 0.250 mm diameter electrodes lies in between the 0.127 and 1mm diameter electrodes, and are approximately 2x the 0.127 mm V_a_, matching theoretical expectations.

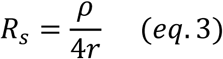

In comparison to V_IPP_ and V_a_ measurements during VTs, which closely match theoretical expectations, the measurements of cathodic and anodic polarization on the electrodes across input impedances are much more complex. This is not unexpected, as in addition to the polarization at the electrode interface having both high frequency and low frequency components, it is also heavily dependent on the surface state of the electrode [32,44,45].

#### 3.3.2 Apparent Qinj Decreased with Sufficient Input Impedance

**Figure 5(a**) shows the impact of the larger observed ΔV_IPP_ highlighted in **Figure 4** on calculated values of E_mc_ and E_ma_ which are used to determine Q_inj_. **Figure 5(b)** shows the mean representative voltage transients collected at the apparent Q_inj_ across electrodes of decreasing surface area without DC mitigation, which can occur at different current densities depending on the electrode size and input impedance of the VT measurement system. At 1 MΩ of input impedance, the apparent Q_inj_ is disproportionately higher for smaller electrodes (215 ± 3.31, 250 ± 15.6 µC/cm^2^ for 0.250 mm and 0.127 mm diameters, respectively), and the polarization direction leading to the potential limit breach varies by electrode size – E_ma_ for the 1mm and 0.250 mm diameter electrodes versus E_mc_ for the 0.127 mm electrodes. As input impedance increases, the breach consistently occurs at E_ma_ across all electrode sizes and at much lower charge densities. This pattern underscores how measurement errors driven by a high R_dc_/Z_in_ ratio can lead to an overestimation of the Q_inj_ of platinum electrodes due to an underestimation of ΔV_IPP_ at the electrode interface which is depicted in **Figure 5(c)**. Significant differences in experimentally determined Q_inj_ were observed between the 1 MΩ/10 MΩ and 10 TΩ groups but not between 10GΩ and 10TΩ groups. Therefore, 10 GΩ was deemed a sufficient input impedance for the voltage transients performed with DC mitigation.

**Figure 5:**
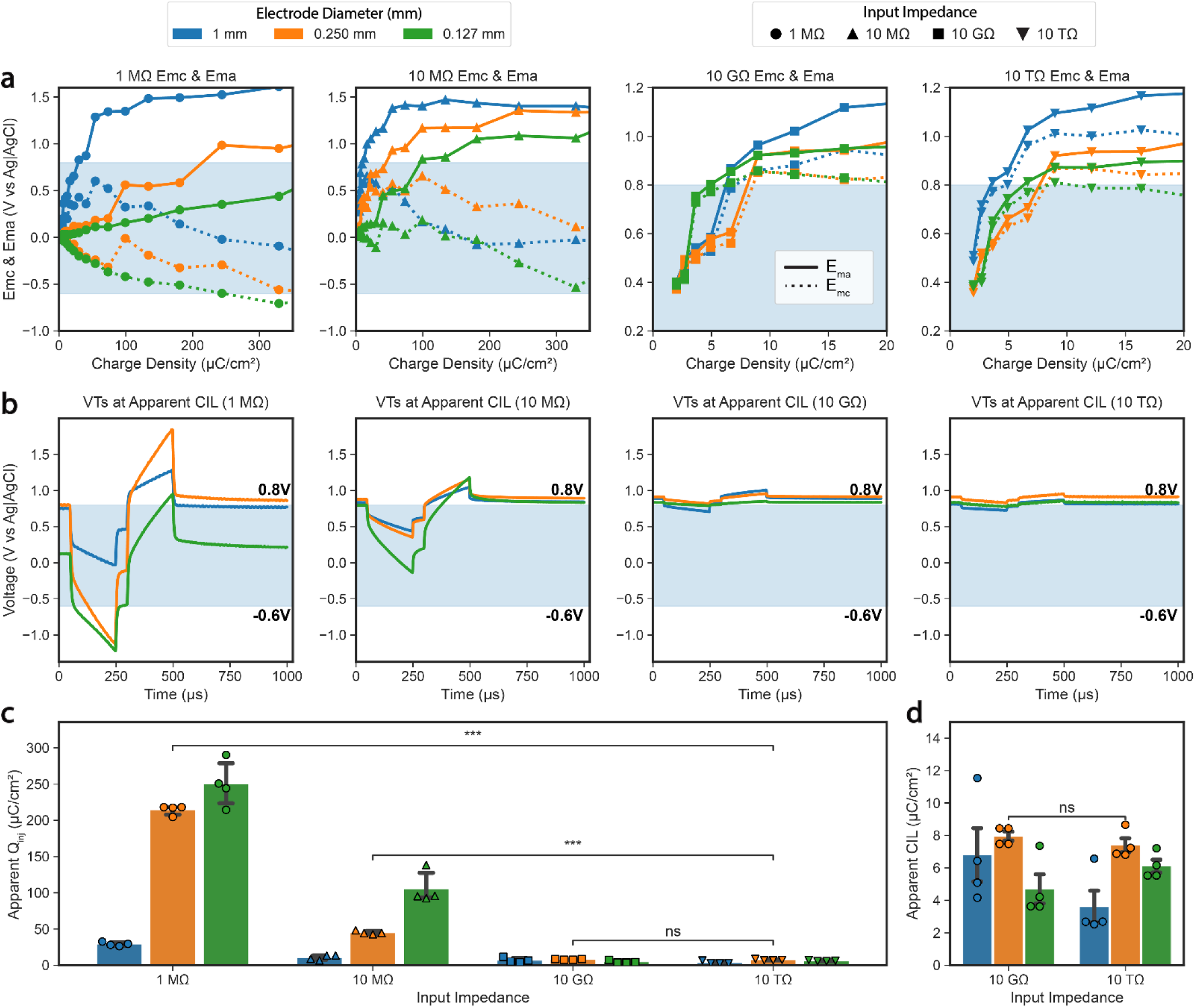
**Traditional methods for calculating charge injection limits (Q_inj_) are susceptible to changes in the ratio of DC electrode impedance to the measurement device input impedance (R_dc_/Z_in_). (**a) Maximum anodic and cathodic polarization (E_mc,_ E_ma_) calculated for the last pulse across input impedances and sizes. (b) Representative mean voltage transient response of the last pulse applied at the charge injection limit (Q_inj_) across sizes. (c-d) Corresponding Q_inj_ calculated from data shown in b. All data in panel c-d are represented as mean ± SEM. *** for p < 0.001. These data were collected without DC mitigation.

### 3.4 Impact of DC Mitigation Strategy on Apparent Charge Injection Limit

Two common methods used to mitigate stimulator leakage currents during stimulation with imbalanced or balanced biphasic pulses are the use of an inline capacitor (CC) and/or the use of an inline capacitor while shorting the cathode/anode in between pulses (CC+ES) [11,18,46–50]. The impedance of a capacitor is inversely proportional to frequency, such that an ideal capacitor functionally has an infinite impedance at DC. Consequently, the inline ‘blocking’ capacitor is used to block stimulator leakage currents from reaching the electrodes. However, a capacitor allows current flow until the voltage across the capacitor equals the voltage of the source (i.e. the stimulator). As shown in **Figure 6** below, when just an inline capacitor is used without shorting the cathode and anode in between pulses, a growing DC offset at the working electrode is still evident over the course of the first ∼400 pulses (**See** **Figure 6(b)**, 1st, 2nd panel; **Figure 6(c)**). Simultaneously, the voltage across the counter electrode blocking capacitor increases linearly due to the unmitigated leakage current of the Keithley current source (**Figure 6(d)**). Around pulse 425, the voltage across the capacitor plateaus and one phase of the biphasic waveform is increasingly ‘cut off’, colloquially known as ‘clipping’ or ‘railing’, resulting in biphasic current waveforms that are no longer charge balanced (See **Figure 6(b)**, 3rd and 4th panel; **Figure 6(d)**). This phenomenon occurs because the voltage of the system (i.e. the sum of the voltages across both blocking capacitors, and both the working and counter electrodes) has reached the 20V compliance limit of the stimulator. Note, at this point V_IPP_ begins to decrease, ultimately settling at a new equilibrium point around ∼0.6V (See **Figure 6(d)**, 4th panel) putatively due to the compromised charge balance of the biphasic current waveform counteracting the stimulator leakage current.

**Figure 6:**
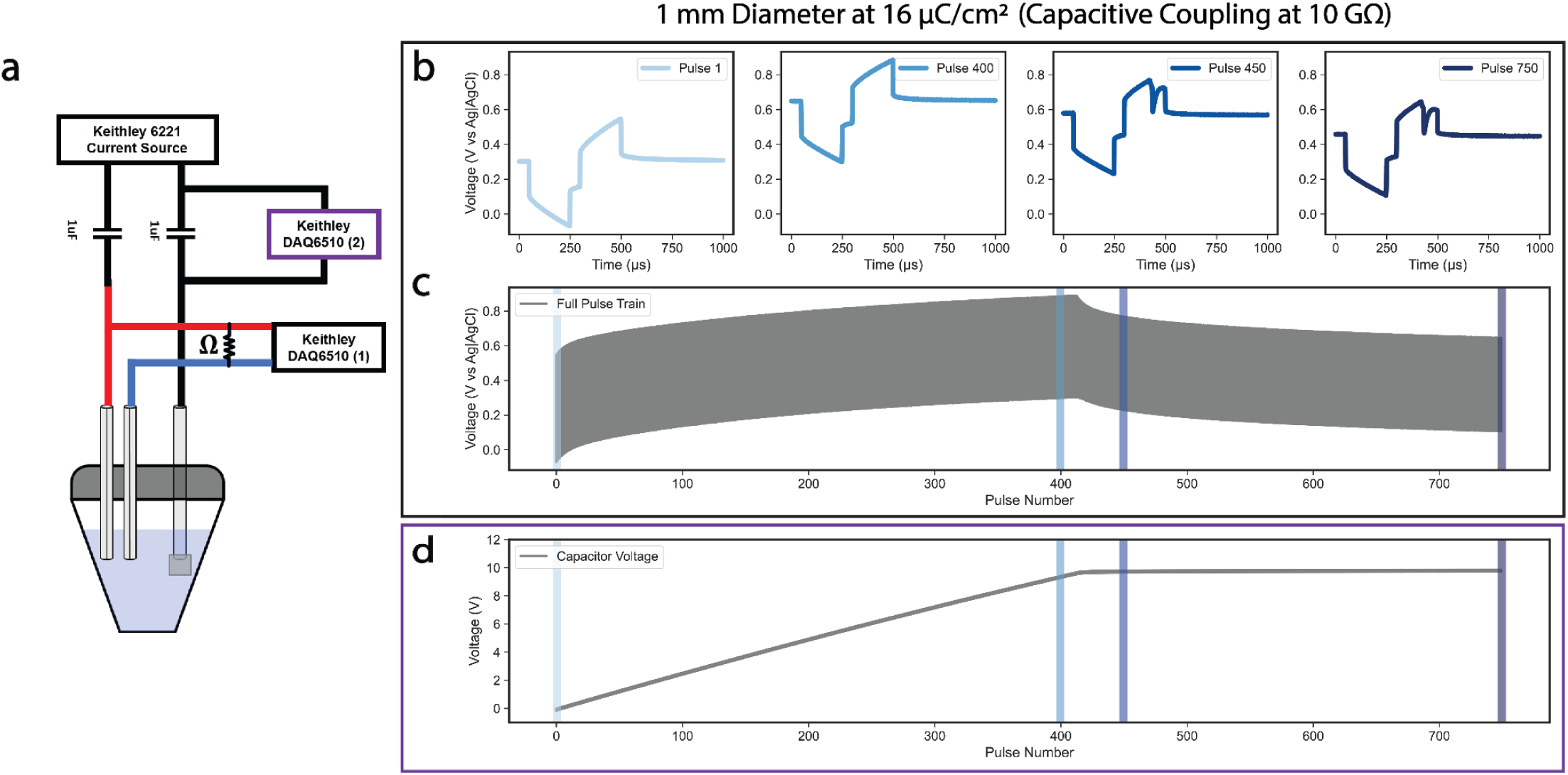
Capacitive coupling alone does not natively improve charge balance of constant current stimulators. (A) Specific experimental configuration used to collect the data displayed in panels b-d. (B) Representative voltage transient pulses (1, 400, 450, 750) collected at a 1 mm diameter electrode over the course of a pulse train at ∼16µC/cm^2^ during the capacitive coupling (CC) condition. (C) Raw voltage plotted against pulse number with colored regions illustrating regions of the pulse train that are represented in panel a. (D) Voltage across the counter electrode blocking capacitor as a function of pulse number.

For this reason, many clinical implantable pulse generators (IPGs) use both an inline capacitor and short behind the electrodes in between pulses, to bleed off charge that has accumulated on the capacitor and prevent clipping/railing of the biphasic pulse. Unfortunately, many stimulators used for preclinical animal research and benchtop testing of electrodes either lack a DC leakage current mitigation strategy or, if capacitively coupled, do not short the cathode/anode between biphasic pulses. This is exacerbated by the fact that measurement systems with insufficient input impedances are often used to measure the output waveform, which as shown previously can dramatically underestimate changes in V_IPP_.

**Figure 7(a) and 7(b)** contrasts the lack of a DC mitigation strategy with CC+ES on a representative example of a single voltage transient trial at 54 µC/cm^2^ with 10 GΩ of input impedance. In this example, all 750 voltage transient pulses for each condition are plotted on the same axis for each electrode size. The corresponding insets represent the ΔV_IPP_ observed over the course of the pulse train. It is evident from these plots that a significant reduction in ΔV_IPP_ was observed with CC+ES. **Figure 8** further elaborates on this observation by comparing the average ΔV_IPP_ as a function of the charge densities applied to each electrode with and without proper DC mitigation. Across the charge densities tested, CC+ES resulted in a reduction of average ΔV_IPP_ ≥ 88% as measured at an appropriate input impedance.

**Figure 7:**
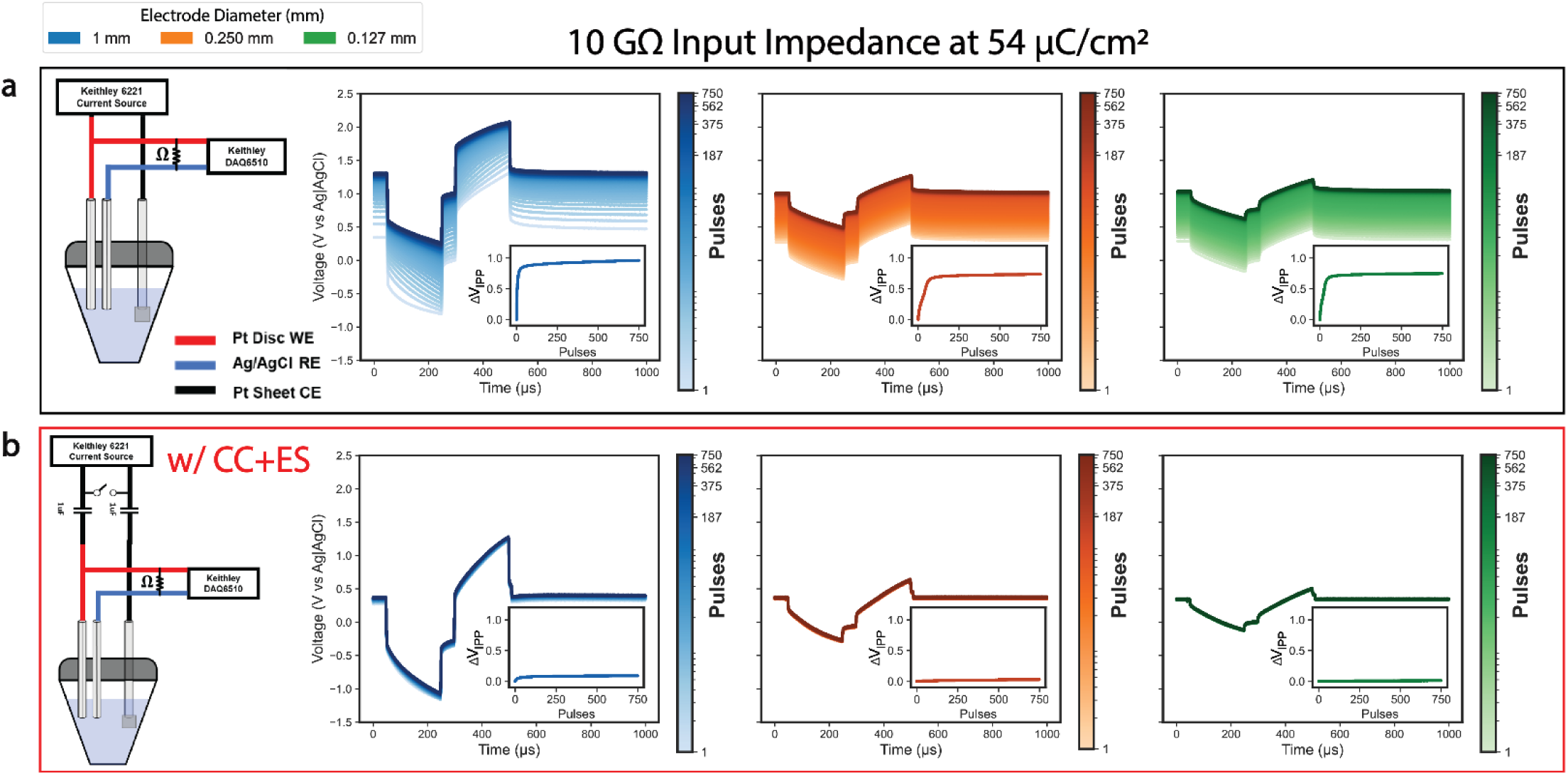
Representative examples showing that capacitive coupling with electrode shorting (CC+ES) greatly reduces the DC offset (ΔV_IPP_) observed during voltage transients. Representative voltage transients collected at 54µC/cm^2^ and 10 GΩ of input impedance without (a) and with (b) custom DC mitigation strategy across sizes. Insets of each plot represent the change in ΔV_IPP_ observed for that trial.

**Figure 8:**
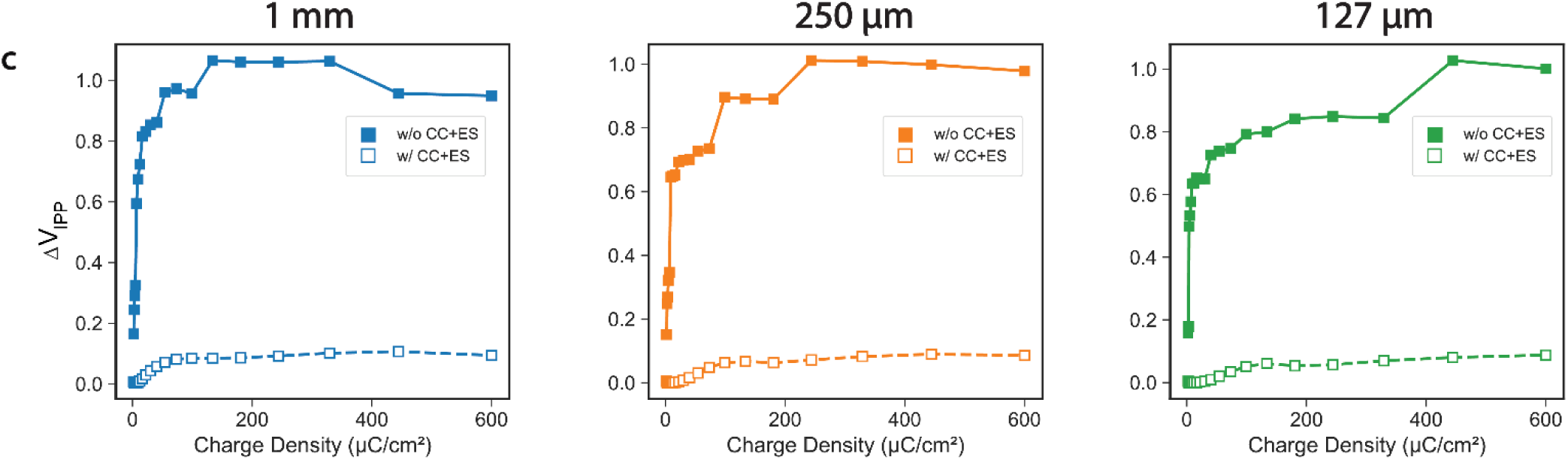
Capacitive coupling with electrode shorting (CC+ES) greatly reduces the DC offset (ΔV_IPP_) observed during voltage transients across charge densities. Average ΔV_IPP_ vs charge density collected with and without custom DC mitigation strategy at 10 GΩ of input impedance across sizes. Note, the ΔV_IPP_ plots for the trials without DC mitigation are transformed from the data presented in Figure 4(B) at 10 GΩ.

The results from the DC mitigation trials across input impedances are presented in **Figure 9**. **Figure 9(a)** displays the Emc and Ema measurements calculated on the 750^th^ pulse across electrode sizes and input impedances. **Figure 9(b)** shows the average representative voltage transients obtained at the apparent Qinj. **Figure 9(c)** presents the charge injection limits calculated across input impedances and electrode sizes. Significant differences in Q_inj_ were observed between the 1 MΩ/10 MΩ and 10 GΩ input impedance Q_inj_ measurements showing that even when a measurement device input impedance still has an impact on experimentally determined Qinj even when an appropriately charge balanced stimulator is used.

**Figure 9:**
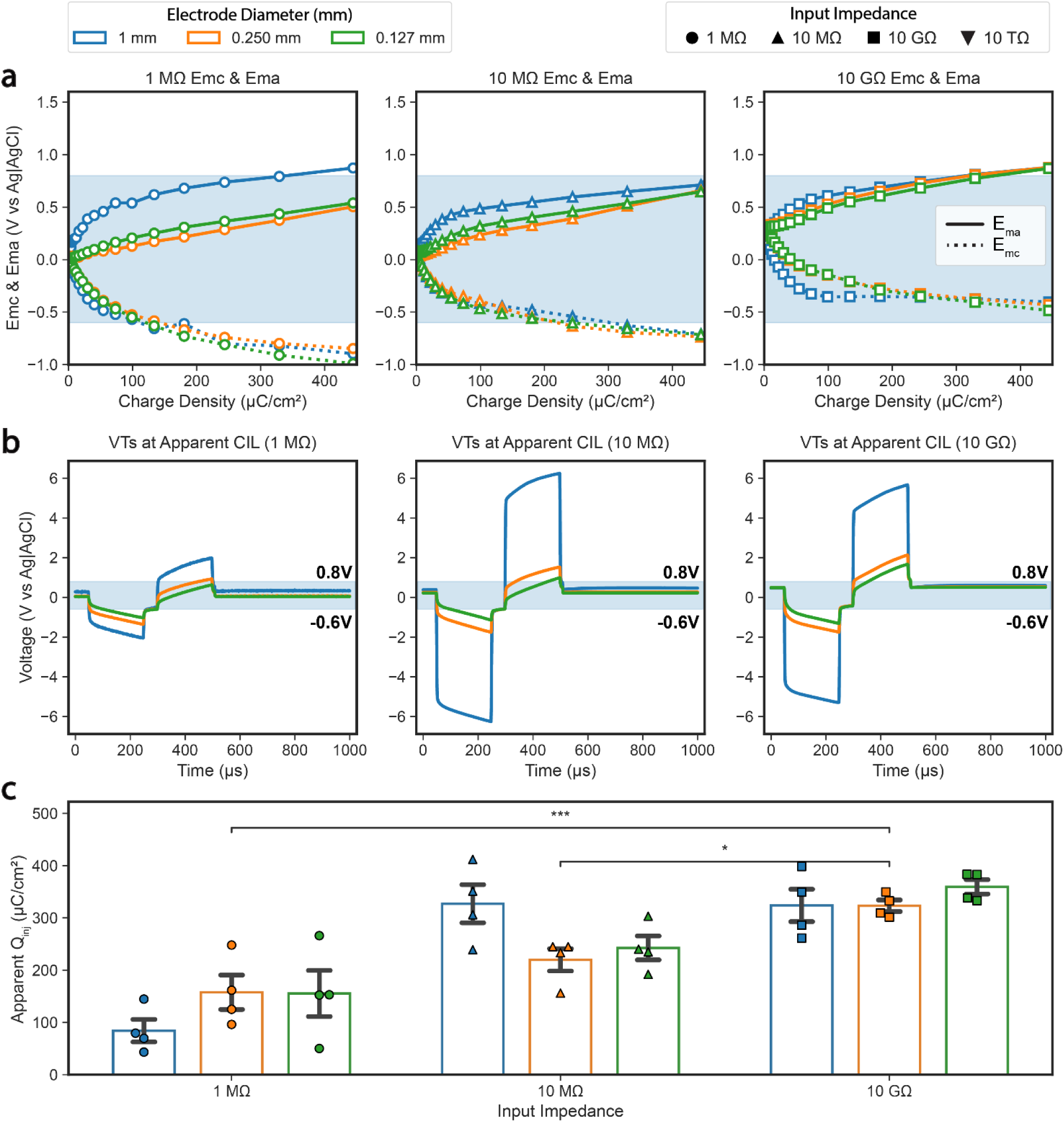
**Traditional methods for assessing charge injection limits (Qinj) are also dependent on DC mitigation strategy (CC+ES). (**a) Maximum anodic and cathodic polarization (E_mc,_ E_ma_) calculated for the last pulse across input impedances and sizes. (b) Representative mean voltage transient response of the last pulse applied at the charge injection limit (Q_inj_) across sizes. (c) Corresponding Q_inj_ calculated from data shown in b. All data in panel c are represented as mean ± SEM.* for p < 0.05 and *** for p < 0.001. These data were collected with DC mitigation.

As discussed in **Section 3.3**, the VT measurements performed with sufficient input impedance and without DC mitigation greatly reduced the calculated Q_inj_ at platinum electrodes using traditional methods. This reduction is due to unmitigated stimulator leakage currents leading to larger observed changes in V_IPP_. However, when these leakage currents are properly mitigated, this alters Q_inj_. **Figure 10(a)** plots the Q_inj_ as a function of R_dc_/Z_in_ across the input impedances, electrode sizes and DC mitigation strategies tested while also contextualizing these results in reference to the historical 50-150 µC/cm^2^ limits which are shaded in green. Note, the low input impedance, DC mitigated trials consistently produced Q_inj_ that were in line with the values derived by Robblee, et al. Without proper DC mitigation, as R_dc_/Z_in_ decreases, so does the calculated Q_inj_ (i.e. 250 ± 15.6, 105 ± 11.0, and 4.71 ± 0.894 µC/cm^2^ at 1 MΩ, 10 MΩ, 10 GΩ, respectively for the 0.127 mm diameter electrodes); however, in this study, an inverse relationship was observed with proper DC mitigation (i.e. 155 ± 44.1, 243 ± 23.0, 359 ± 13.8 µC/cm^2^ at 1 MΩ, 10 MΩ, 10 GΩ, respectively for the 0.127 mm diameter electrodes). This suggests that with appropriate measurement instrumentation in addition to a properly charge balanced stimulator, appreciably larger charge injection limits can be safely applied [46]. **Figure 10(b)** represents an alternative grouping of these results that demonstrates the average effect of DC mitigation on the calculated Q_inj_ across different input impedances. ***These data stress that both the ratio of the DC electrode impedance to the measurement device input impedance and the DC mitigation strategy deployed have a significant impact on Q_inj_*.**

**Figure 10:**
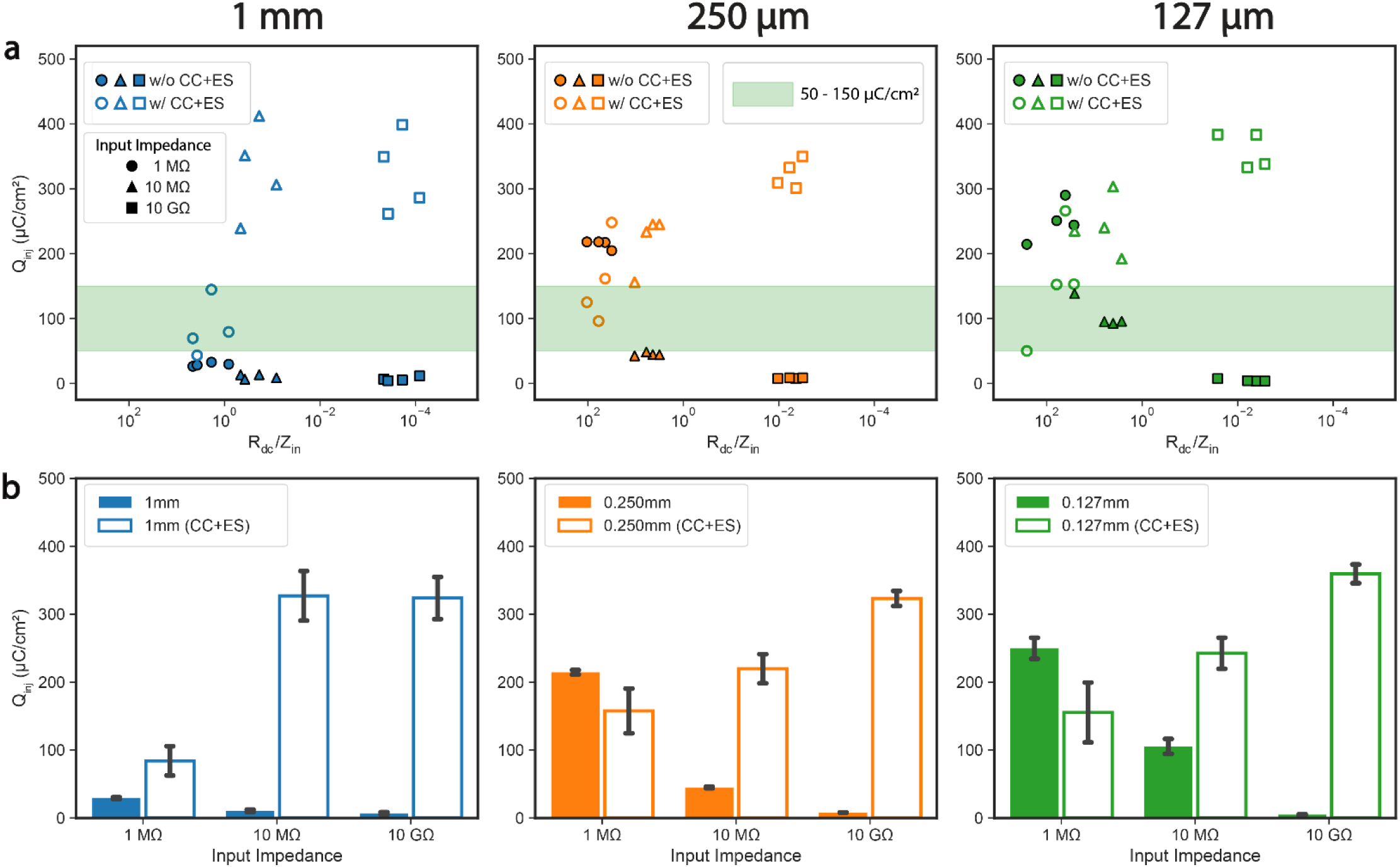
**Experimentally derived charge injection limits (Q_inj_) can differ widely depending on both R_dc_/Z_in_ and DC mitigation strategy**[51]. (a) R_dc_/Z_in_ (as determined by EIS around the open circuit potential (OCP)) vs charge injection limit for each sample across sizes, input impedances with and without DC mitigation. The highlighted region in green represents the 50-150 µC/cm^2^ limits determined by Robblee, 1990 [12]. (b) Charge injection limits across sizes, with and without DC mitigation, grouped by input impedance. Solid bars in panel b are identical to the charge injection limits represented in Figure 5(c). Hollow bars in panel b are identical to the charge injection limits represented in Figure 9(c). All data in panel b are represented as mean ± SEM.

## 4. Discussion

To safely stimulate neural tissue, charge can be delivered through either capacitive or faradaic charge transfer mechanisms. Capacitive charge transfer is generally considered the safest way to inject charge for neural stimulation. Conversely, charge passed through faradaic mechanisms can be unsafe if the reactions are not completely reversed leading to the formation of toxic byproducts and/or corrosion of the electrode. In practice, all electrodes have a limited amount of charge that can be passed exclusively through safe reactions. A fundamental and unresolved challenge in designing stimulation electrodes is determining stimulation limits that enable sufficient charge injection to achieve therapeutic effect but prevent reactions that are unsafe to tissue or damage the electrode.

Voltage transients (VTs) are routinely used to assess what reactions are available for faradaic charge transfer and how much current can be passed before the onset of unsafe reactions. In this paper, we demonstrate that the accuracy of voltage transient measurements can be impacted by stimulation and measurement set-ups commonly used in literature. These data stress the importance of characterizing both the stimulator set-up as well as the measurement system to enable consistent and accurate determination of Q_inj_ from VTs. Based on these results, in the following discussion we comment on the importance of accurately monitoring DC electrode potentials and mitigating stimulator leakage currents at neural stimulators. Additionally, we provide a comprehensive table detailing reporting strategies enabling reproducible VTs for the characterization of novel electrode materials, the development of neural stimulators, and the evaluation of complete neuromodulation systems.

### 4.1 Effect of Measurement Instrumentation on DC Electrode Potential

An ideal voltage measurement would have infinite input impedance, resulting in an accurate measurement of the DC electrode potential that is not subject to probe loading introduced by the measurement device [52]. In practice, oscilloscopes, typically with input impedances of 1-10 MΩs, are commonly used to measure electrode voltage while conducting VTs. This is much lower than the >10 TΩ input impedance for typical electrochemical instrumentation (i.e. potentiostats). When the input impedance of the measurement system is too low relative to the DC impedance of the electrode, changes in V_IPP_ are poorly resolved, leading to incorrect assessments of when the electrode reaches the limits of water electrolysis. ***This undermines the core premise of voltage transient analysis, which is to infer the onset of irreversible faradaic processes from the observed electrode interfacial polarization***. Without proper tracking of V_IPP_, particularly in two-electrode configurations or systems lacking sufficient measurement device input impedance, it becomes difficult to accurately determine the electrode’s potential relative to the desired electrochemical window. As a result, it may not be possible to accurately infer if the measured electrode potential could lead to the formation of unsafe reaction products or degradation of the electrode. As such, accurate measurement and reporting of V_IPP_ is a prerequisite for determining parameters of safe stimulation across studies [36,44,45,53].

### 4.2 DC Electrode Potential as an Indicator of Stability during Voltage Transients (Charge Balance vs Electrochemical Balance)

The DC potential of an electrode reflects the balance of available reactants, reaction products in the surrounding electrolyte as well as the electrochemical state of the surface [44,54]. For Pt electrodes, there exists distinct polarization regions in which different faradaic pathways dominate, each with varying charge transfer capabilities. While electrochemical safety is often determined by electrode polarizations leading to the electrolysis of water (i.e. -0.6V and +0.8V vs Ag|AgCl), at lower levels of polarization, other reactions occur that can lead to the loss of electrode material and/or the formation of toxic byproducts. For example, the formation of platinum chloride begins at ∼0.5V vs Ag|AgCl, well within the nominal water window, and has been linked to accelerated Pt dissolution and neural toxicity [55–57].

Additionally, the reactions that occur within the water window are not perfectly reversible. While charge balanced pulses are most commonly used to stimulate neural tissue, for a given pulse a portion of that charge goes into irreversible electrochemical processes. As a result, irreversible reactions consume reactants and accumulate reaction products, perturbing the local electrode environment over time, resulting in a residual DC offset that reflects the underlying reaction asymmetry. This distinction between charge balance (equally injected/withdrawn current) and electrochemical balance (no net chemical change at the interface) is critical [17]. Different electrode materials rely on unique faradaic pathways to support charge transfer, which inherently differ in their reaction and mass transfer kinetics. Several studies report that deliberately ‘breaking’ charge balance, via small biases or asymmetric pulses, can reduce dissolution at some materials by driving the interface toward a more electrochemically favorable operating point [58–60]. However, as demonstrated in **Section 3.4**, an improperly charge-balanced stimulator can produce electrode polarization shifts that are unrelated to the intrinsic electrochemical behavior of the electrode. Therefore, before using DC electrode potential as an indicator of electrochemical stability, it is necessary to verify that instrumentation-related sources of DC electrode polarization have been minimized.

To disentangle instrumentation-driven effects from true electrochemical behavior we applied the same stimulation amplitudes and DC mitigation strategies used in-vitro to a simple dummy load (**Supplementary Figure 5a**). These measurements serve two purposes: 1) To validate the impact of the DC mitigation strategies on the stimulator leakage current and 2) To demonstrate the differences in stimulator leakage current specifications across common preclinical stimulators (Keithley 6221 vs TDT Subject Interface) despite delivering the same charge-balanced waveform (**Supplemental Figure 5b-e**). In contrast to the dummy load, which exhibited zero leakage current using our mitigation strategy (**Supplementary Figure 5b**), the in-vitro measurements exhibited small but consistent shifts in V_IPP_ **(Figure 8)**. Their presence under truly charge-balanced stimulation indicate that the faradaic reactions supporting charge transfer are not perfectly reversible.

Taken together, our findings support the growing view that a properly measured DC offset, once the instrumentation-related confounds are rectified, provides a meaningful readout of the electrochemical stability at the electrode interface [19].

### 4.3 Avoiding Confounds: Practical Experimental Considerations and Reporting Strategies

*Rigorous reporting of the specifications of the instrumentation (measurement device input impedance (Z_in_), stimulator leakage currents, DC mitigation strategies, stimulator output impedance (Z_out_)) used to determine the Q_inj_ for a particular electrode is critical for the interpretation of electrochemical test results across studies and materials* [36]. To understand the prevalence of studies evaluating electrochemical performance which may be impacted by inappropriate measurement instrumentation or improperly mitigated DC leakage currents, we performed a methodological survey as described in **Supplementary Methods**. Of the 77 studies matching the inclusion criteria, 73 reported the use of an oscilloscope as the VT measurement instrumentation. Of those studies, 3 reported the probe input impedance. Additionally, while 72/77 studies reported the stimulator used, only 23 reported the use of a DC mitigation strategy. Critically, 22/77 studies indicated the use of a ‘custom stimulator’ without reporting stimulator specifications (**Supplementary Table 5**).

Our data demonstrates that differences in instrumentation can create widely varying reports of Q_inj_. Consistent with that, our methodological survey conducted herein found reports of Q_inj_ ranging from 150 – 5570 µC/cm^2^ for platinum electrodes [61], and 24 – 23000 µC/cm^2^ for TiN electrodes [62–64]. We suspect that the lack of methodological consistency has contributed significantly to this variation in Q_inj_ observed across the literature.

#### 4.3.1 Practical Considerations for Reproducible and Consistent Assessment of Electrode Materials using Voltage Transients

Preventing corrosion and reaction byproduct accumulation are crucial to ensuring the longevity and safety of neuromodulation devices. This presents a larger issue at microelectrodes versus macroelectrodes due to smaller margin of error before critical failure, especially for chronically implanted active electrodes. The data presented herein provide a plausible explanation as to why premature failure of TFMEAs is so common. To avoid unwanted electrochemical reactions, it is crucial to accurately measure the DC electrode potential to understand the electrochemical reactions being utilized to pass charge[53]. Additionally, as mentioned in **Section 4.2**, instrumentation confounds like stimulator leakage currents should be minimized to elucidate the true electrochemical behavior of the material under investigation. While platinum was the only material characterized in this study, the results are relevant and can be extrapolated to other electrode materials. For example, electrodes and electrode coatings that are faradaic in nature (i.e. IrOx) will have a lower R_ct_, meaning that these electrodes require a lower Z_in_ to properly measure the DC electrode potential. In contrast, electrode materials and coatings that are more capacitive in nature (i.e. TiN) will have an inherently higher R_ct_ requiring a higher Z_in_. Electrode surface area and fractal geometry will also influence these electrode parameters. A faradaic electrode with a larger electroactive surface area will ultimately lower R_ct_ and thus reduce R_dc_/Z_in_ [65,66]. Studies characterizing novel electrode designs and materials for stimulation should use **Table 2** to carefully consider whether their instrumentation provides an accurate representation of the Q_inj_ of the material.

**Table 2:**
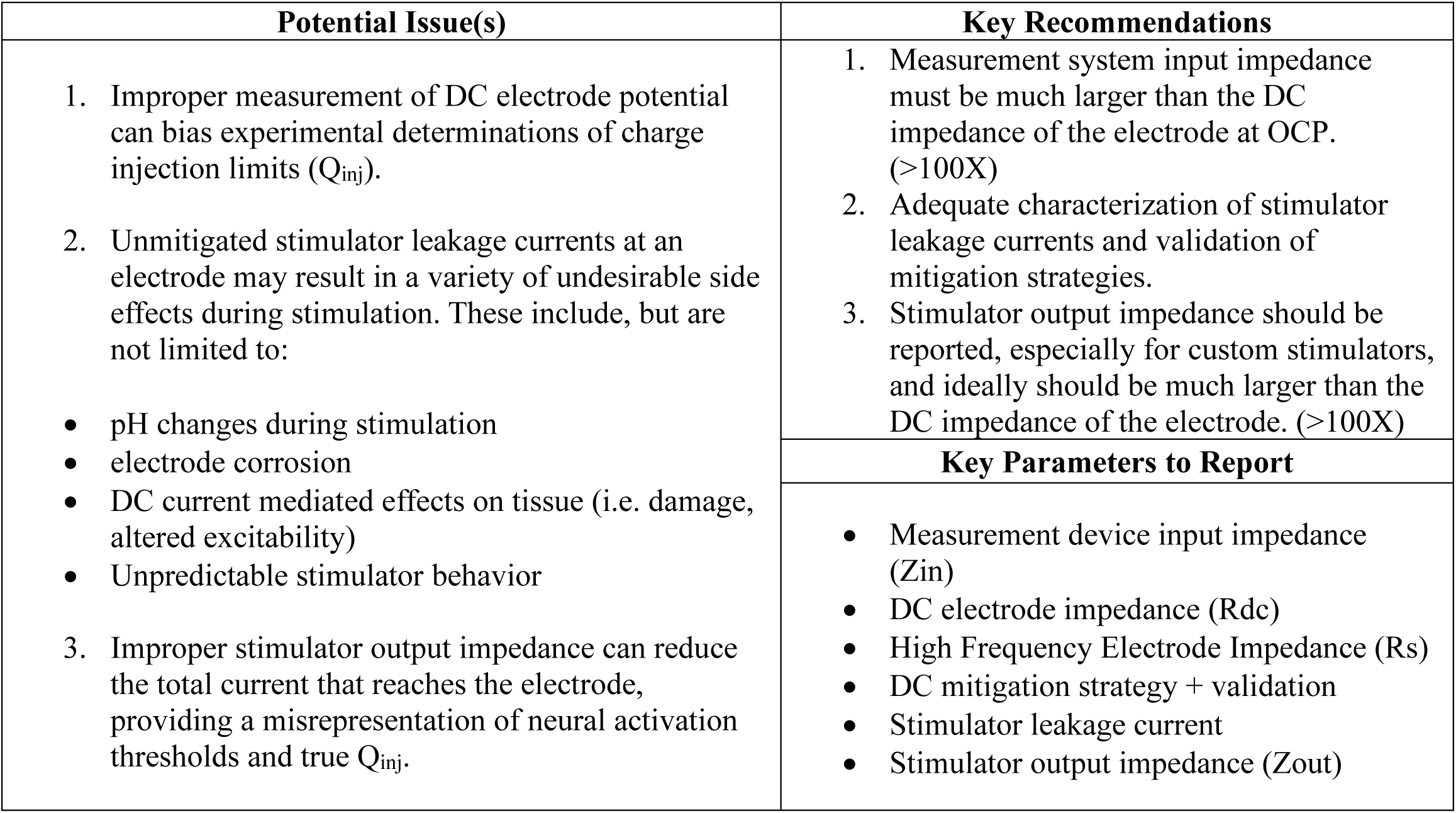
Key Issues and Reporting Recommendations for Functional Neuromodulation, Neural Interface Materials and Neurostimulator Development Studies

| Potential Issue(s) | Key Recommendations |
| --- | --- |
| Improper measurement of DC electrode potential can bias experimental determinations of charge injection limits ($Q_{inj}$).<br>Unmitigated stimulator leakage currents at an electrode may result in a variety of undesirable side effects during stimulation. These include, but are not limited to:<br>pH changes during stimulation<br>electrode corrosion<br>DC current mediated effects on tissue (i.e. damage, altered excitability)<br>Unpredictable stimulator behavior<br>Improper stimulator output impedance can reduce the total current that reaches the electrode, providing a misrepresentation of neural activation thresholds and true $Q_{inj}$. | Measurement system input impedance must be much larger than the DC impedance of the electrode at OCP. ($>100X$)<br>Adequate characterization of stimulator leakage currents and validation of mitigation strategies.<br>Stimulator output impedance should be reported, especially for custom stimulators, and ideally should be much larger than the DC impedance of the electrode. ($>100X$) |
|  | Key Parameters to Report |
|  | Measurement device input impedance ($Z_{in}$)<br>DC electrode impedance ($R_{dc}$)<br>High Frequency Electrode Impedance ($R_s$)<br>DC mitigation strategy + validation<br>Stimulator leakage current<br>Stimulator output impedance ($Z_{out}$) |

#### 4.3.2 Practical Considerations and Implications for Neural Stimulation in Functional Neuromodulation Experiments

In vitro charge injection limits are often used to justify that the stimulation parameters applied during in-vivo experiments fall within electrochemically safe boundaries. Findings suggest that unmitigated DC leakage currents may impact both in-vivo functional responses and long-term safety outcomes (i.e. electrode failure, tissue damage). Inadequate DC mitigation can lead to unintended charge accumulation at the electrode interface, resulting in local pH shifts, sustained DC electric fields or other effects that may alter neuronal excitability and tissue health [21,24–27]. Additionally, improper DC mitigation can cause unpredictable stimulator behavior, such as waveform truncation, which may go unnoticed (**Section 3.4**). These consequences are particularly relevant for small electrodes, like those commonly used in preclinical rodent models, where small stimulator leakage currents can result in large shifts in DC electrode potential. Consequently, there may be cases within the literature where Q_inj_ and, by extension, the safety margins of a given neuromodulation system or material, are a misrepresentation of the true functional and electrochemical performance. Future studies should ensure stimulator leakage currents are appropriately mitigated for a given electrode **(Table 2)**. Additionally, future work should critically assess how instrumentation-related errors may have influenced historically reported functional and safety related outcomes.

#### 4.3.3 Practical Considerations for Neurostimulator Development

The use of DC blocking capacitors on stimulator output stages is considered good practice both clinically and preclinically to prevent prolonged DC accumulation at an electrode. However, as demonstrated in **Section 3.4**, simply adding capacitors does not result in a charge balanced stimulator that effectively eliminates ΔV_IPP_. In fact, infrequent discharge of the DC blocking capacitors can result in decreased stimulator voltage compliance, or the accumulation of significant and potentially harmful offset voltages [67,68]. As noted in this study, once the stimulator reaches compliance, waveform asymmetry and truncation can occur, undermining charge balance and leading to unpredictable or undesirable electrochemical behavior at the electrode-electrolyte interface. (**Figure 6**).

The DC mitigation strategy proposed in this study required the use of large DC blocking capacitors on the stimulator output. This approach becomes impractical with multichannel, high density implantable devices. As a result, several alternative DC mitigation strategies that have been proposed in recent years to reduce their physical footprint and cost [69–71]. To rigorously validate the effectiveness of these alternative strategies in vitro, it is imperative to use appropriate measurement device input impedances to properly measure DC accumulation, particularly for TFMEAs due to their susceptibility to corrosion-related failures and higher impedance.

In addition to high input impedance measurement instrumentation, it is critical to verify that the stimulator has adequate output impedance for the intended application. Sufficient output impedance ensures that the intended current reaches the electrode interface and is not lost through parasitic pathways (**Table 2**).

### 4.5 Limitations & Future Directions

When interpreting the results of this study, several limitations should be considered. The platinum electrodes used in this study were hand-fabricated using materials from various manufacturers. Although care was taken to standardize the surface condition, variability in surface morphology and potential contamination remain possible sources of error. Additionally, the absence of pre-trial control over V_IPP_ and electrolyte reactant concentrations may have contributed to some of the variability in Q_inj_ across trials. In some voltage transient trials, it was observed that V_IPP_ did not completely return to pre-trial levels, particularly when large changes in V_IPP_ occurred; likely reflecting persistent changes in the local electrode environment. Future work could improve reproducibility by incorporating potentiostatic control of V_IPP_ prior to stimulation.

This study used the platinum electrolysis window as a reference framework for assessing Q_inj_, consistent with many previous reports. However, it is well established that there are corrosion/noxious reactions at platinum electrodes within the water window (i.e. formation of reactive oxygen species, platinum oxidation) [45,56,72,73]. Since electrochemical reactivity and failure mechanisms are material-specific, future studies should assess whether the electrolysis window remains a valid framework for evaluating Q_inj_ across emerging neuromodulation materials.

This study also tested a very specific set of parameters (i.e. pulse widths, amplitudes, frequencies) on the observed DC offset. Future experiments should explore the effects of waveform parameterization on DC offset magnitude and evaluate how sustained DC offsets affect electrode performance across different neuromodulation materials. Understanding these material-specific failure modes is critical for assessing long term safety and reliability of neuromodulation devices. In parallel, future studies should also characterize the effectiveness of DC mitigation strategies across these materials to determine whether existing approaches provide adequate protection or require adaptation for specific electrode compositions and geometries.

While we demonstrated the implications of electrode size on probe loading, the smallest electrodes used in this study were considerably larger than what is typically considered a microelectrode (< 2 × 10^-5^ cm^2^). With these electrodes we found that 10GΩ of input impedance was sufficient to properly resolve the DC electrode potential, but additional research should be performed to understand the measurement instrumentation requirements TFMEAs as a function of electrode size and intercontact distance. The electrodes tested in this study were isolated single electrodes and are not representative of the parasitics that would be experienced with a high-density electrode array with electrode contacts of equivalent surface area. Therefore, the results presented herein represent a ‘best-case scenario’ with regards to potential parasitic pathways that exist.

Lastly, neural electrode safety is a multifactorial problem that cannot be completely assessed via in-vitro performance tests. In-vitro tests like voltage transients allow for rapid comparisons between devices to be readily made, but as demonstrated by our results, these comparisons must be contextualized in reference to the instrumentation used [72]. It is important to note that in-vitro tests are not a measure of safety alone but give an idea for what types of electrochemical reactions may occur at the electrode interface during a particular therapy. There are several additional confounds that exist in the in-vivo electrode environment due to its dynamic nature that are difficult to replicate in-vitro. Aspects like the protein adsorption and the foreign body response can drastically change the electrode interface including electrode properties like impedance, corrosion resistance and Q_inj_ [73,74]. Doses that cause clinically significant tissue damage will be location and tissue dependent and should be evaluated on a case-by-case basis. However, the data presented herein do have practical implications in vivo, as the DC observed in-vitro is typically much lower than in-vivo [26]. With several reports indicating that chronic stimulator leakage currents in excess of 100 nA are associated with higher levels of tissue damage; by properly monitoring and mitigating them we can appreciably improve preclinical and clinical device safety while also maximizing longevity [22,26,75].

## 5. Conclusion

This study examined the role of measurement device input impedance and DC mitigation strategies in accurately determining Q_inj_ using voltage transients. Our findings demonstrate that insufficient input impedance leads to significant underestimation of shifts in V_IPP_. These errors can obscure the onset of irreversible faradaic processes, misrepresenting electrode safety and compromising the validity of electrochemical testing.

Unmitigated stimulator leakage currents also pose serious risks to long-term device performance and tissue safety. Sustained V_IPP_ drift has been shown to accelerate corrosion, alter local pH and promote the accumulation of toxic reaction byproducts. These risks are especially pronounced in microelectrodes, which have reduced tolerance to electrochemical imbalance before critical failures. Unmitigated/uncharacterized DC can also alter electrophysiological outcomes, potentially causing poor reproducibility of key physiological findings across studies [24,27,28].

As such, we emphasize the need for standardized reporting of instrumentation parameters including measurement device input impedance, stimulator leakage currents, stimulator output impedance and implemented DC mitigation strategies when reporting Q_inj_. Electrode properties like R_ct_ and R_s_ should also be disclosed to establish the influence of measurement configuration on observed V_IPP_ values. **The accuracy and reproducibility of Q_inj_ measurements depend not only on the electrode material but on the entire electrochemical system and methodology used to interrogate it.**

## Data Availability

All data that support the findings of this study are included within the article (and any supplementary files). Any additional data are available from the corresponding author upon reasonable request.

## Acknowledgements

We sincerely thank Dr. Justin Williams, Dr. Aaron Suminski, Dr. Brandon Coventry, and Dr. Harry Finklea for reviewing the manuscript and providing incredibly insightful feedback. We also would like to express our gratitude to the members of the Wisconsin Institute for Translational Neuroengineering (WITNe) for their unwavering support of this work.

