## Supplementary Material for "Systematic Errors in DC Offset Measurement and Mitigation at Neurostimulation Electrodes: Causes, Implications and Solutions"

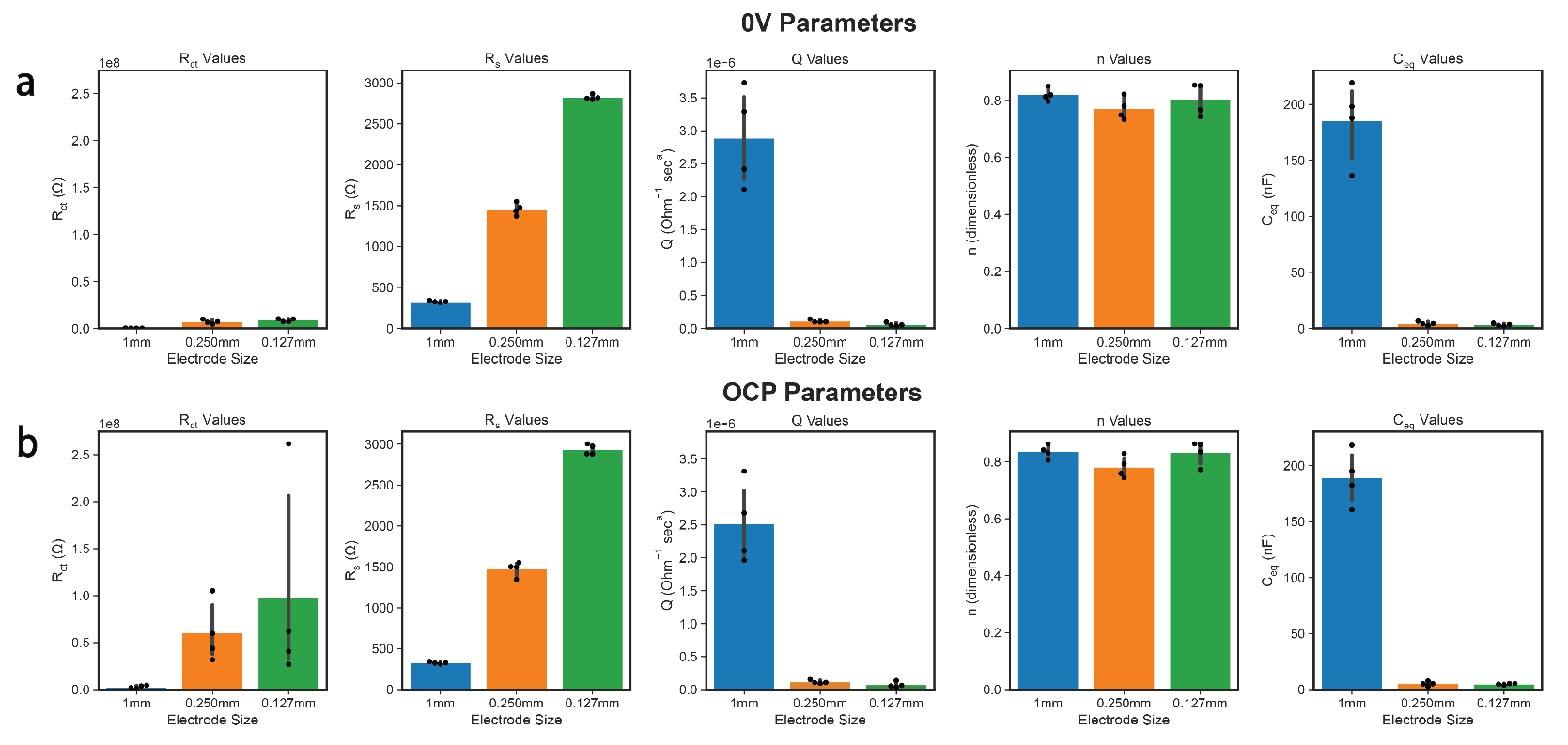

### **Supplementary Information**

**Supplementary Figure 1:** **Visualization of Randles fit values across electrode sizes.** Bar plot of Randles circuit fit parameters determined from EIS spectra collected at 0V relative to Ag|AgCl (a) and OCP (b).

**Supplementary Table 1:** Raw model fit parameters determined from Randles circuit fit of EIS spectra collected at 0V relative to Ag|AgCl across the 12 samples tested.

| **Sample ID** | **Electrode Size** | **Rct (Ω)** | **Rs (Ω)** | **Q (Ohm^-1^ sec^a^)** | **n** |
| --- | --- | --- | --- | --- | --- |
| P1 | 1 mm | 2.24E+05 | 3.25E+02 | 3.29E-06 | 8.13E-01 |
| P2 | 1 mm | 2.34E+05 | 3.26E+02 | 2.42E-06 | 8.51E-01 |
| P3 | 1 mm | 2.02E+05 | 3.12E+02 | 3.73E-06 | 7.97E-01 |
| P4 | 1 mm | 3.05E+05 | 3.41E+02 | 2.11E-06 | 8.20E-01 |
| QPI1 | 0.250 mm | 7.10E+06 | 1.55E+03 | 1.42E-07 | 7.49E-01 |
| QPI2 | 0.250 mm | 6.42E+06 | 1.44E+03 | 9.66E-08 | 7.80E-01 |
| QPI3 | 0.250 mm | 4.60E+06 | 1.48E+03 | 9.79E-08 | 8.22E-01 |
| QPI4 | 0.250 mm | 9.81E+06 | 1.37E+03 | 9.69E-08 | 7.33E-01 |
| EPI1 | 0.127 mm | 1.02E+07 | 2.87E+03 | 5.24E-08 | 7.66E-01 |
| EPI2 | 0.127 mm | 1.01E+07 | 2.80E+03 | 9.45E-08 | 7.44E-01 |
| EPI3 | 0.127 mm | 7.28E+06 | 2.82E+03 | 4.94E-08 | 8.52E-01 |
| EPI4 | 0.127 mm | 7.18E+06 | 2.81E+03 | 3.59E-08 | 8.54E-01 |

**Supplementary Table 2:** Raw model fit parameters determined from Randles circuit fit of EIS spectra collected at the OCP relative to Ag|AgCl across the 12 samples tested.

| **Sample ID** | **Electrode Size** | **Rct (Ω)** | **Rs (Ω)** | **Q (Ohm^-1^ sec^a^)** | **n** |
| --- | --- | --- | --- | --- | --- |
| P1 | 1 mm | 4.62E+06 | 3.42E+02 | 1.97E-06 | 8.41E-01 |
| P2 | 1 mm | 1.88E+06 | 3.23E+02 | 2.11E-06 | 8.62E-01 |
| P3 | 1 mm | 8.20E+05 | 3.12E+02 | 3.31E-06 | 8.05E-01 |
| P4 | 1 mm | 3.76E+06 | 3.26E+02 | 2.68E-06 | 8.31E-01 |
| QPI1 | 0.250 mm | 1.05E+08 | 1.50E+03 | 1.06E-07 | 8.29E-01 |
| QPI2 | 0.250 mm | 4.37E+07 | 1.50E+03 | 1.08E-07 | 7.93E-01 |
| QPI3 | 0.250 mm | 3.18E+07 | 1.35E+03 | 9.53E-08 | 7.43E-01 |
| QPI4 | 0.250 mm | 5.98E+07 | 1.55E+03 | 1.52E-07 | 7.59E-01 |
| EPI1 | 0.127 mm | 2.62E+08 | 2.88E+03 | 5.35E-08 | 8.60E-01 |
| EPI2 | 0.127 mm | 4.07E+07 | 3.00E+03 | 6.27E-08 | 8.35E-01 |
| EPI3 | 0.127 mm | 2.69E+07 | 2.97E+03 | 3.84E-08 | 8.63E-01 |
| EPI4 | 0.127 mm | 6.21E+07 | 2.88E+03 | 1.37E-07 | 7.72E-01 |

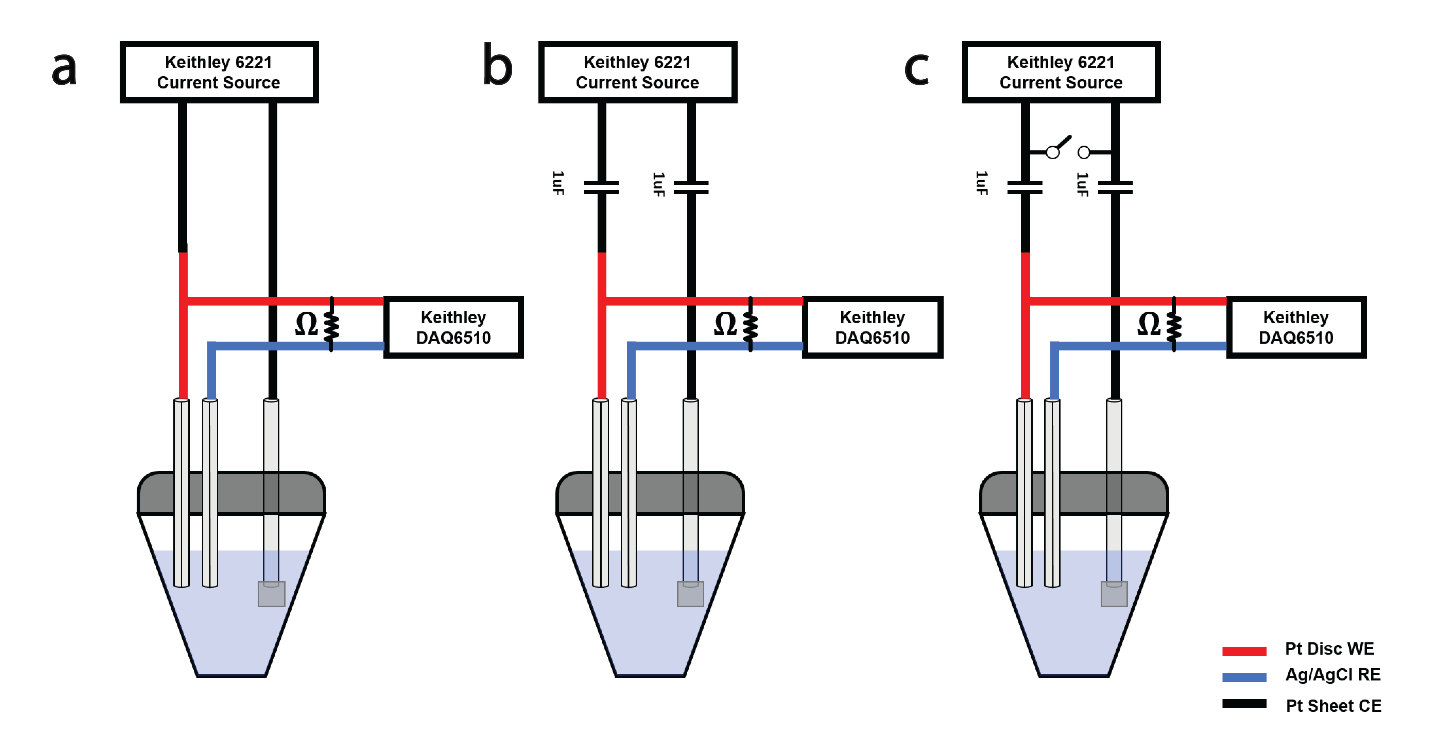

**Supplementary Figure 2: Stimulator/measurement configurations for the voltage transient DC mitigation strategies compared in this study.** (a) Lack of DC mitigation strategy, (b) Capacitive coupling only (CC), and (c) Capacitive coupling with electrode shorting (CC+ES)

**Supplementary Table 3:** Charge densities and resultant current amplitudes applied in this study across electrode sizes.

| **Charge Densities (µC/cm^2^)** | **Applied Current Amplitudes (µA)** | | |
| --- | --- | --- | --- |
|  | **1 mm** | **0.250 mm** | **0.127 mm** |
| 2.00 | 78.54 | 4.91 | 1.27 |
| 2.70 | 106.04 | 6.63 | 1.71 |
| 3.65 | 143.17 | 8.95 | 2.32 |
| 4.92 | 193.29 | 12.08 | 3.13 |
| 6.65 | 260.97 | 16.31 | 4.22 |
| 8.97 | 352.34 | 22.02 | 5.70 |
| 12.11 | 475.71 | 29.73 | 7.69 |
| 16.36 | 642.27 | 40.14 | 10.39 |
| 22.08 | 867.14 | 54.20 | 14.02 |
| 29.81 | 1170.75 | 73.17 | 18.93 |
| 40.25 | 1580.66 | 98.79 | 25.56 |
| 54.34 | 2134.09 | 133.38 | 34.51 |
| 73.37 | 2881.30 | 180.08 | 46.59 |
| 99.06 | 3890.12 | 243.13 | 62.90 |
| 133.74 | 5252.16 | 328.26 | 84.93 |
| 180.57 | 7091.09 | 443.19 | 114.66 |
| 243.80 | 9573.87 | 598.36 | 154.81 |
| 329.16 | 12925.95 | 807.86 | 209.01 |
| 444.40 | 17451.68 | 1090.72 | 282.20 |
| 600.00 | 23562.00 | 1472.61 | 381.00 |

**Supplementary Methods:**

**DC Mitigation Circuitry Characterization**

To ensure that the parasitics of the DC mitigation circuitry did not appreciably impact the experimental results presented in this manuscript, we measured the impedance spectra between traces of an unpopulated DC mitigation PCB. Impedance spectra were collected using an Autolab (PGSTAT12). This allowed us to determine the differential trace impedances of the PCB. Success was determined by differential trace impedances comparable to the off-impedance of the DG417L switch (~10 GΩ).

**Rs Calculation**

Validation of the methodology used to calculate the charge injection limit (Q_inj_) involved back-calculating the value of Rs using the subtracted access voltage (V_a_) and the applied current. The resultant value was then compared to the values of Rs extracted from the Randles circuit fit of the EIS spectra.

**Methodological Survey**

A methodological survey was conducted on 4/15/26 to identify studies reporting charge injection limits/capacities derived from voltage transient measurements across the neuromodulation literature. A unified Boolean search strategy was deployed using the following search string input into Google Scholar: ("voltage transient" OR "voltage excursion")  AND  ("charge injection limit" OR "charge injection capacity") AND ("oscilloscope"). This search revealed 165 results. Articles were screened by three independent reviewers to confirm that they met the inclusion criteria (**see below**) resulting in a total of 77 unique results **(Supplementary Table 3)**. Subsequently, the relevant parameters (**see below**) were ascertained from each qualifying record. Notably, database-specific differences impacted record retrieval. Instrumentation details such as the use of oscilloscopes are rarely reported in titles, abstracts, or indexed keywords, and are instead typically confined to full-text methods sections. As a result, the unified Boolean search strategy with the identical string input across Scopus and PubMed yielded zero search results and thus was not included.[1–77]

***It is important to note that simply using an oscilloscope or 2-electrode setup does not inherently mean that the charge injection limits are inaccurate. This methodological survey is strictly meant to highlight the common use of these tools across the neuromodulation literature and to motivate the relevance of this manuscript.***

Inclusion criteria:

- Conduct voltage transients as part of electrochemical characterization.
- Used an oscilloscope or comparable input impedance device as measurement device for voltage transients.
- Reported in-vitro charge injection limits/capacity for device/material.
- Electrode setup could be 3-electrode or 2-electrode.
- Preprints, dissertations, and review papers were excluded.

Extracted Parameters:

- Recording Instrument
- Recording Impedance
- Stimulator
- DC Mitigation Strategy
- Charge Injection Limit (µC/cm^2^)
- Material
- Electrode GSA (cm^2^)
- Electrode Configuration

**Supplementary Table 4:** Raw values of Rs extracted from EIS spectra and voltage transient Q_inj_ calculation method as presented in **Supplementary Figure 3**.

| **Electrode Diameter (mm)** | **Sample ID** | **Rs EIS** | **1 MΩ** | | **10 MΩ** | | **10 GΩ** | |
| --- | --- | --- | --- | --- | --- | --- | --- | --- |
|  |  |  | **Rs VT** | **Rs VT (CC+ES)** | **Rs VT** | **Rs VT (CC+ES)** | **Rs VT** | **Rs VT (CC+ES)** |
| 1 mm | P1 | 3.25E+02 | 3.63E+02 | 3.86E+02 | 3.36E+02 | 4.14E+02 | 3.29E+02 | 3.07E+02 |
| 1 mm | P2 | 3.26E+02 | 3.23E+02 | 3.44E+02 | 3.54E+02 | 3.86E+02 | 3.36E+02 | 3.37E+02 |
| 1 mm | P3 | 3.12E+02 | 3.30E+02 | 3.81E+02 | 3.43E+02 | 4.30E+02 | 3.53E+02 | 3.44E+02 |
| 1 mm | P4 | 3.41E+02 | 3.22E+02 | 4.19E+02 | 3.42E+02 | 3.99E+02 | 3.42E+02 | 3.33E+02 |
| 0.250 mm | QPI1 | 1.55E+03 | 1.40E+03 | 1.85E+03 | 1.57E+03 | 1.87E+03 | 1.55E+03 | 1.42E+03 |
| 0.250 mm | QPI2 | 1.44E+03 | 1.48E+03 | 1.65E+03 | 1.60E+03 | 1.66E+03 | 1.55E+03 | 1.42E+03 |
| 0.250 mm | QPI3 | 1.48E+03 | 1.29E+03 | 1.49E+03 | 1.58E+03 | 1.66E+03 | 1.49E+03 | 1.35E+03 |
| 0.250 mm | QPI4 | 1.37E+03 | 1.43E+03 | 1.76E+03 | 1.53E+03 | 1.70E+03 | 1.49E+03 | 1.45E+03 |
| 0.127 mm | EPI1 | 2.87E+03 | 3.26E+03 | 3.71E+03 | 3.10E+03 | 3.12E+03 | 3.21E+03 | 3.17E+03 |
| 0.127 mm | EPI2 | 2.80E+03 | 2.85E+03 | 3.33E+03 | 3.56E+03 | 2.93E+03 | 3.39E+03 | 2.80E+03 |
| 0.127 mm | EPI3 | 2.82E+03 | 3.18E+03 | 3.34E+03 | 3.36E+03 | 2.97E+03 | 3.27E+03 | 2.79E+03 |
| 0.127 mm | EPI4 | 2.81E+03 | 3.15E+03 | 3.06E+03 | 3.32E+03 | 2.94E+03 | 3.32E+03 | 2.84E+03 |

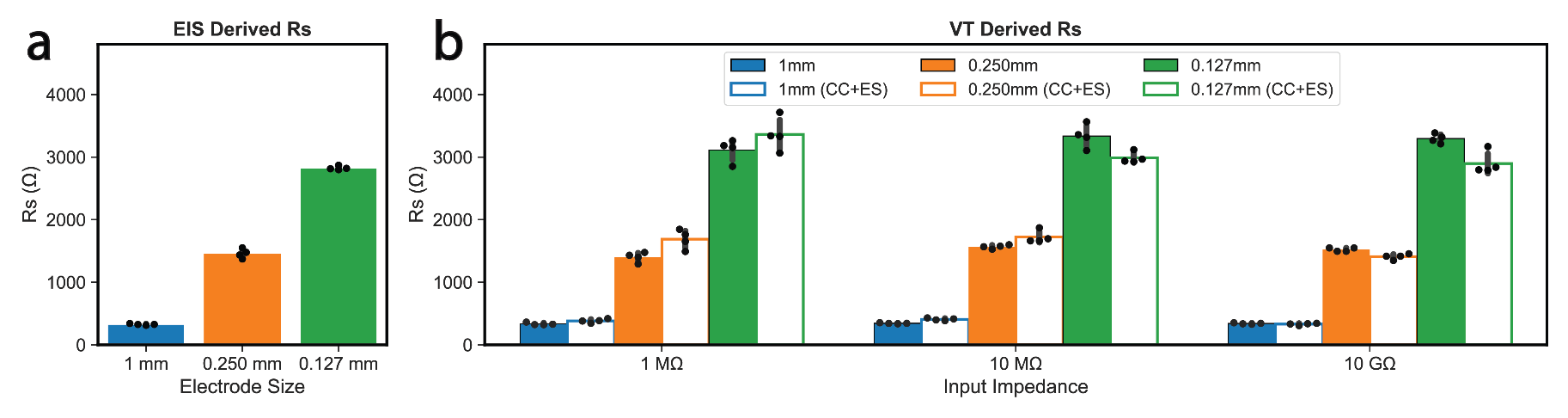

**Supplementary Figure 3: Validation of methodology used to calculate charge injection limits (Q_inj_).** Comparison of R_s_ values attained using electrochemical impedance spectroscopy (EIS) Randles circuit fit (a) and the voltage transient charge injection limit (Q_inj_) calculation method (b).

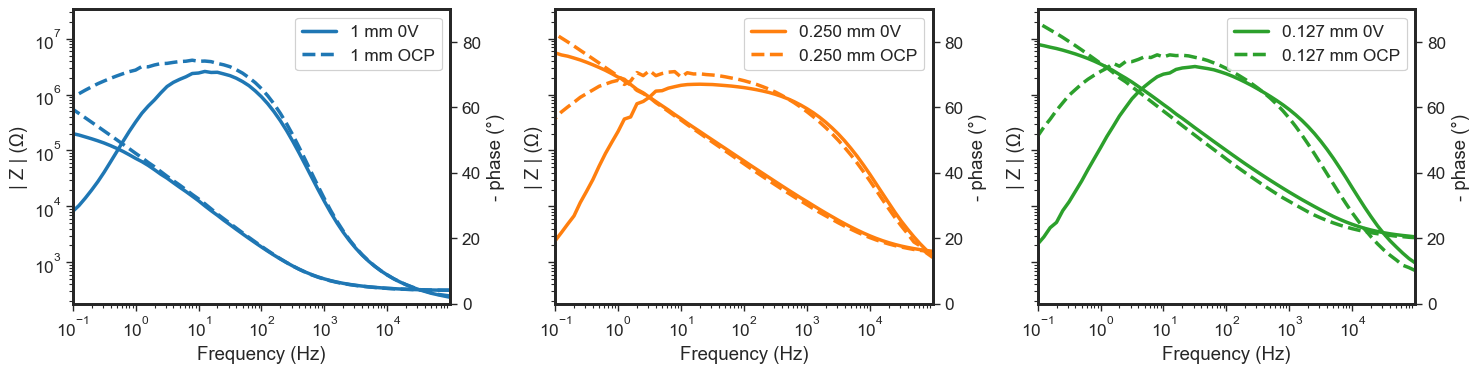

**Supplementary Figure 4: Comparison of OCP vs 0V vs Ag|AgCl EIS spectra across electrode sizes.** This figure demonstrates the effect of bias voltage on the EIS spectra. Note the difference in low frequency behavior as a result of collecting EIS around OCP rather than 0V vs Ag|AgCl.

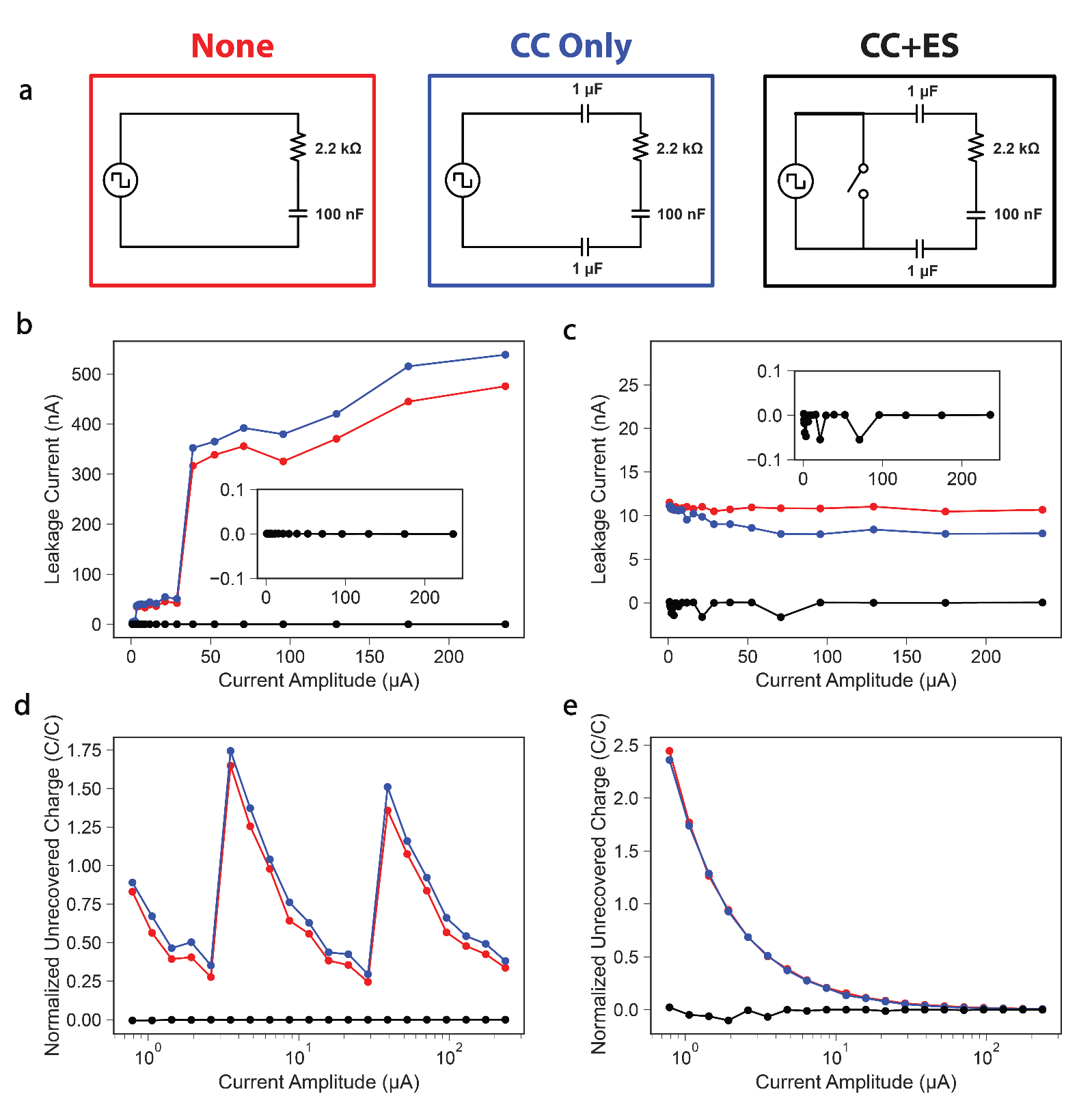
**Supplementary Figure 5: Stimulator leakage currents vary widely across preclinical stimulators.** (a) DC mitigation strategies used to measure unrecovered charge across a dummy cell. Measured leakage current and unrecovered charge normalized to the charge per phase applied during each current amplitude for the Keithley 6221 Current Source (b & d) and the Tucker Davis Technologies Subject Interface (c & e). The current amplitudes represented in this figure were the same as the currents applied to the 0.250 mm diameter electrodes.

***Supplementary References***

[62]	Woodington B J, Lei J, Carnicer A, Güemes A, Naegele T E, Hilton S, El S, Trivedi R A, Malliaras G G and Barone D G 2024 Flexible circumferential bioelectronics to enable 360-degree recording and stimulation of the spinal cord *Sci. Adv.*
